# Systematic discovery of Short Linear Motifs decodes calcineurin phosphatase signaling

**DOI:** 10.1101/632547

**Authors:** Callie P. Wigington, Jagoree Roy, Nikhil P. Damle, Vikash K. Yadav, Cecilia Blikstad, Eduard Resch, Cassandra J. Wong, Douglas R. Mackay, Jennifer T. Wang, Izabella Krystkowiak, Devin Bradburn, Eirini Tsekitsidou, Su Hyun Hong, Malika Amyn Kaderali, Shou-Ling Xu, Tim Stearns, Anne-Claude Gingras, Katharine S. Ullman, Ylva Ivarsson, Norman E. Davey, Martha S. Cyert

## Abstract

Short linear motifs (SLiMs) drive dynamic protein-protein interactions essential for signaling, but sequence degeneracy and low binding affinities make them difficult to identify. We harnessed unbiased systematic approaches for SLiM discovery to elucidate the regulatory network of calcineurin (CN)/PP2B, the Ca^2+^-activated phosphatase that recognizes LxVP and PxIxIT motifs. *In vitro* proteome-wide detection of CN-binding peptides, *in vivo* SLiM-dependent proximity labeling, and *in silico* modeling of motif determinants uncovered unanticipated CN interactors, including NOTCH1, which we establish as a CN substrate. Unexpectedly, CN shows SLiM-dependent proximity to centrosomal and nuclear pore complex (NPC) proteins – structures where Ca^2+^ signaling is largely uncharacterized. CN dephosphorylates human and yeast NPC proteins and promotes accumulation of a nuclear transport reporter, suggesting conserved NPC regulation by CN. The CN network assembled here provides a resource to investigate Ca^2+^ and CN signaling and demonstrates synergy between experimental and computational methods, establishing a blueprint for examining SLiM-based networks.

## Introduction

Cellular networks signal through dynamic protein-protein interactions (PPIs) that are often formed by low-affinity docking of short linear interaction motifs (SLiMs) onto globular domains. SLiMs are degenerate 3-10 amino acid long sequences within intrinsically disordered regions that evolve rapidly and harbor many disease-causing mutations (Davey et al., 2017; Uyar et al., 2014). By determining complex formation, modification state, localization, and stability, SLiMs control much of the decision making that directs a protein from translation to degradation (Van Roey et al., 2014), yet many SLiMs remain to be identified (Tompa et al., 2014). Multiple protein phosphatases recognize substrates via specific SLiMs (Brautigan and Shenolikar, 2018; Kataria et al., 2018; Ueki et al., 2019). Thus, rapidly developing methods for SLiM discovery (Davey et al., 2017; Krystkowiak et al., 2018) can provide a systems-level understanding of phosphatases, which has heretofore been limited by challenges in identifying their substrates.

Calcineurin (CN/PP2B/PPP3) is the only Ca^2+^/calmodulin-regulated phosphoserine/threonine-specific protein phosphatase. Although this highly conserved, ubiquitously expressed enzyme critically regulates the cardiovascular, nervous and immune systems, only ~70 proteins are established as CN substrates (Table S5). Because CN dephosphorylates Nuclear Factor of Activated T-cells (NFAT) transcription factors to promote T-cell activation (Jain et al., 1993), CN inhibitors are clinically important immunosuppressants (Liu, 1993). However, these drugs, FK506 (Tacrolimus) and cyclosporin A (CsA), cause many unwanted effects, such as hypertension, diabetes, and seizures by inhibiting CN in non-immune tissues (Roy and Cyert, 2019). Systematic mapping of CN signaling pathways will therefore aid in understanding and potentially ameliorating these negative consequences.

CN, a heterodimer of catalytic (CNA) and regulatory (CNB) subunits, is autoinhibited by sequences in CNA that are displaced when Ca^2+^ binds to CNB and Ca^2+^/calmodulin to CNA (Roy and Cyert, 2019). Conserved docking surfaces on CN bind PxIxIT and LxVP SLiMs, whose positioning within substrates varies (Figure 1A) (Roy and Cyert, 2019). These motifs play distinct roles during dephosphorylation: PxIxIT targets CN to substrates and regulators by binding CN under both basal and signaling conditions (Li et al., 2007). In contrast, LxVP binds to a site that is accessible only in active CN, and is thought to orient phosphorylated residues toward the active site (Grigoriu et al., 2013; Li et al., 2016). Mutating PxIxIT or LxVP motifs in substrates or their cognate binding surfaces on CN disrupts dephosphorylation, and CN inhibitors (CsA, FK506, and the viral A238L protein) act by blocking SLiM binding (Grigoriu et al., 2013). Although these SLiMs have been characterized at the structural level (Grigoriu et al., 2013; Li et al., 2004; Li et al., 2007; Sheftic et al., 2016), the small number of validated instances in humans has limited understanding of their binding determinants and hampered global identification.

**Figure 1:**
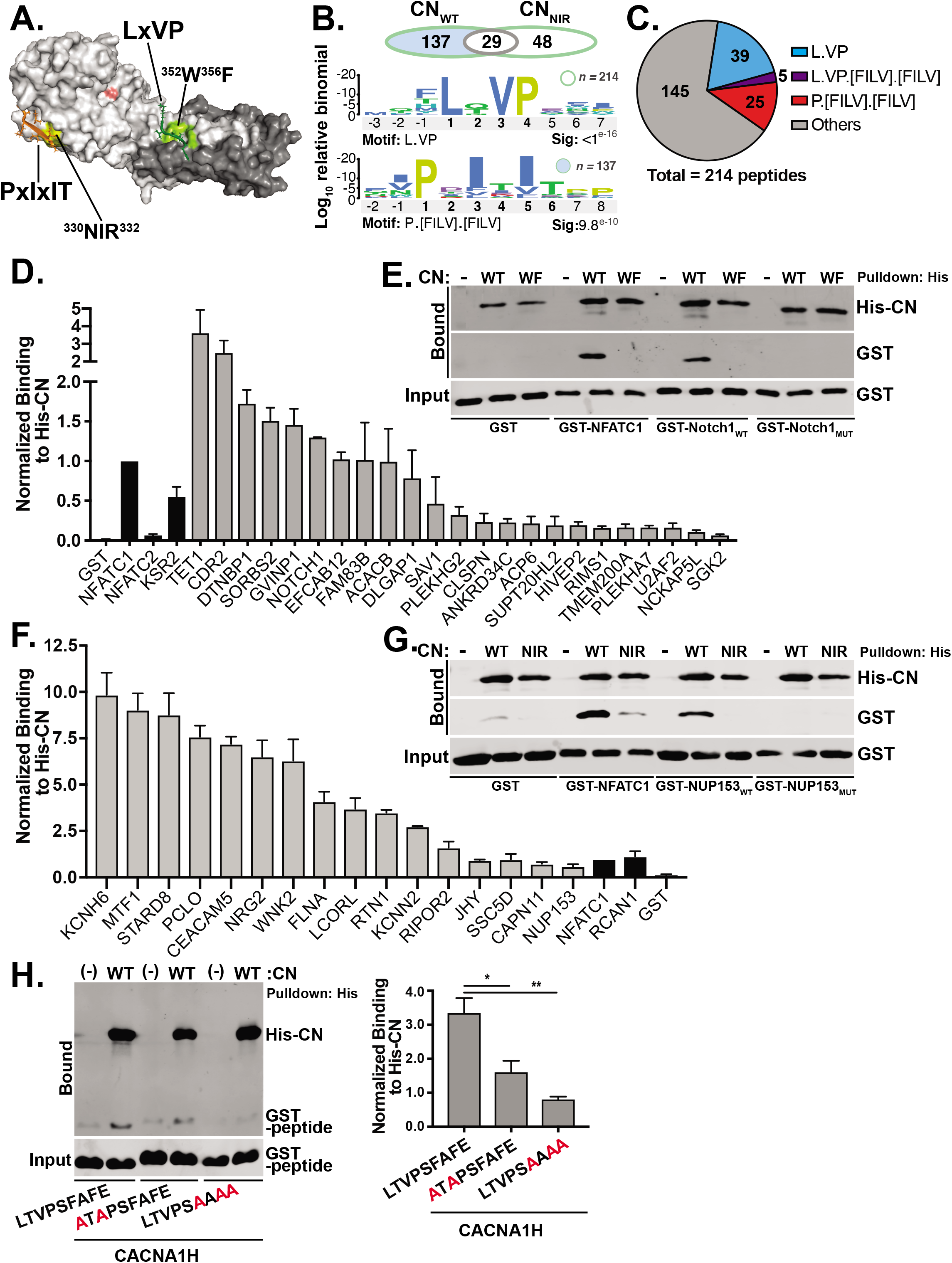
ProP-PD selections with CN discover human PxIxIT and LxVP sequences. **A**. Structure of the CNA/CNB heterodimer (A α = silver, catalytic center = pink, B = grey) complexed with PxIxIT (orange) and LxVP (green) motifs (PDB: 4F0Z). Residues crucial for PxIxIT (^330^NIR^332^: yellow) and LxVP (^352^W,^356^F: lime) docking are shown. **B**. Motif discovery: L.VP was enriched in total Pro-PD peptides (214, green outline); P.[FILV].[FILV] was enriched in peptides selected by CN_WT_ only (137, blue fill). **C**. Motif distribution among ProP-PD peptides. **D**. Copurification of GST-tagged LxVP peptides with His-CN, relative to the LxVP from NFATC1. Data are mean ± SEM; n=3. Known LxVPs (black bars), novel LxVPs (gray bars). In all cases, binding is significantly greater than to GST alone (p<0.01). **E**. Co-purification of GST-tagged LxVP peptides from NFATC1 and NOTCH1 ICD (NOTCH1_WT_: NTPSHQ**L**Q**VP**EHPFLT or NOTCH1_MUT_: NTPSHQ**<u>A</u>**Q**<u>AA</u>**EHPFLT) with His-CN (CN_WT_ or CN_WF_). **F**. Co-purification of GST-PxIxIT peptides with His-CN relative to the PxIxIT from NFATC1. Data are mean ± SEM; n=3. In all cases p<0.01, and bar coloring scheme as in **D**. **G**. Co-purification of GST-tagged PxIxIT peptides from NFATC1 and NUP153 (NUP153WT: L**P**T**F**N**FS**SPEITTSSP or NUP153_MUT_: L**<u>A</u>**T**<u>A</u>**NF**<u>A</u>**SPEITTSSP) with His-CN (CN_WT_ or CN_NIR_). **H**. Co-purification of GST-tagged LxVPxIxIT peptide from CACNA1H (WT or mutants as indicated) with His-CN. Data are mean ± SEM; n=3. *: p<0.05, **: p<0.01. See also Figures S1, S2 and Table S1.

In this work, we discovered unexpected functions and substrates for CN by systematically identifying CN-binding SLiMs in the human proteome. Proteomic peptide phage-display revealed PxIxIT and LxVP sequences in 39 proteins whose association with CN is unexplored, including the NOTCH1 intracellular domain (NICD), which we establish as a CN substrate. Furthermore, by leveraging this expanded collection of CN-binding SLiM sequences we created tools for proteome-wide identification of CN interactors *in silico.* In complementary studies using proximity-dependent biotinylation (BioID) to probe SLiM-dependent associations *in vivo,* we uncovered surprising CN proximities to centrosomal and nuclear pore complex (NPC) proteins. Finally, we demonstrated a conserved function for CN at the NPC by showing that it directly dephosphorylates NPC basket proteins from yeast and humans *in vitro* and regulates nucleoporin phosphorylation and import of a nuclear transport reporter *in vivo.* In this work, experimental and computational approaches synergize to transform our understanding of Ca^2+^/CN signaling and establish a blueprint for future investigations of SLiM-based networks.

## Results

### Proteomic Peptide Phage Display (ProP-PD) uncovers human PxIxIT and LxVP sequences

To directly identify CN-binding SLiMs, CN_WT_ and CN_NIR_, a mutant with reduced binding to PxIxIT SLiMs (Li et al., 2004), were used as bait to select a phage library displaying 16-mer peptides from all intrinsically disordered regions of the human proteome (Figure 1A, S1A, S1B). Final selected phage pools were sequenced and analyzed for enriched motifs using SLiMFinder (Edwards et al., 2007), identifying 214 unique peptides that include 8 known CN-binding SLiMs, some of which were represented by a single sequence read (Table S1).

L.VP was the most highly enriched motif with 44 instances (significance: <1e^-16^), including four established LxVPs (Figure 1B, C, Table S1). Following fusion of these sequences to GST, their co-purification with CN_WT_ was measured semi-quantitatively relative to the LxVP from NFATC1 and GST alone, yielding 23 peptides with variable binding to CN_WT_ and reduced binding to CN_WF_, a mutant that compromises LxVP binding (Figures 1A, D, E, S1D, S2A) (Rodriguez et al., 2009). Furthermore, mutating LxVP within a peptide from NOTCH1 reduced its interaction with CN_WT_, confirming that the motif is critical for binding to CN (NOTCH1_MUT_, Figure 1E).

PxIxIT-like motifs were uncovered by analyzing 137 peptides from selections with CN_WT_, but not the PxIxIT binding-defective CN_NIR_ (Figure 1B, C, S1C) (Li et al., 2004). 30 instances of P.[FILV].[FILV] (henceforth referred to as ‘PxIxIT’) were identified, including 4 previously established PxIxITs (significance: <9.76e^-10^) (Table S1). When fused to GST, 16 of these peptides significantly co-purified with CN_WT_, as measured relative to GST and the PxIxIT from NFATC1, and had reduced binding to CN_NIR_ (Figure 1F, G, S2B). These analyses yielded 18 novel CN-binding sequences, as peptides from SSC5D and LCORL each contained a pair of functional PxIxITs (data not shown). Mutating the PxIxIT in a peptide from NUP153 (NUP153_MUT_; Figure 1G), disrupted its co-purification with CN_WT_, confirming that the identified motif is required for CN-binding.

Interestingly, 5 ProP-PD peptides contained both CN-binding motifs in a combined L.VP.[FILV].[FILV] sequence (Table S1). When tested, 4 of these peptides bound to CN_WT_, and one from KCNJ8 showed only LxVP-type binding (Figure S2A, Table S1). In contrast, the CACNA1H peptide displayed both PxIxIT and LxVP characteristics: binding to both CN_NIR_ and CN_WF_ was significantly reduced vs CN_WT_ (Figure S2C), and mutating either LxV or IxIT disrupted CN_WT_-binding (Figure 1H). Similarly, peptides from C16orf74 and CACUL1 exhibited both binding modes (data not shown). These combined sequences are unlikely to engage LxVP and PxIxIT docking surfaces simultaneously, as they are separated by ~60 Å (Grigoriu et al., 2013). However, each SLiM may bind CN independently, as observed for the “shuffle” complexes formed between MAPK and different docking sites on Nhe1 (Hendus-Altenburger et al., 2016).

The remaining 145 ProP-PD peptides (Figure 1C) were likely enriched via non-specific binding during phage selections, as 24 of 26 tested failed to bind CN_WT_ (data not shown, Table S1).

In summary, these analyses tripled known LxVP and PxIxIT sequences from 22 to 67 (Figure 3A, B), uncovered peptides with characteristics of both motifs, and demonstrate the power of unbiased proteome-wide selection methods to discover SLiMs.

### Human SLiMs identify NOTCH1 as a CN substrate

Our analyses uncovered previously unknown SLiMs in the CN substrates Filamin A and KCNJ8 (Table S1), and in 39 unique proteins whose predicted interaction with CN expands the scope of CN signaling in humans (Figure 2A). These include NUP153, a component of the nuclear pore complex (NPC), and NOTCH1, a developmental regulator (Table S1). *In vivo* associations of NUP153 and NOTCH1 proteins with CN were probed via proximity-dependent labeling using the promiscuous biotin ligase, BirA* (BioID), which can detect low-affinity, SLiM-mediated interactions (Gingras et al., 2018; Roux et al., 2012). Consistent with the demonstrated PxIxIT in NUP153, biotinylation of FLAG-NUP153 was reduced with BirA*-CN_NIR_ vs -CN_WT_, and biotinylation of FLAG-NUP153 with a mutated PxIxIT (PxIxIT_MUT_) was reduced with both enzymes (Figure 2B). NOTCH1 encodes a transmembrane receptor that is cleaved after ligand binding to produce an intracellular domain (NICD; Figure 2C) that functions as a transcription factor (Tzoneva and Ferrando, 2012). Consistent with the demonstrated LxVP in NOTCH1, NICD_WT_-FLAG was biotinylated significantly more than NICD-LxVP_MUT_-FLAG, in which the LxVP is mutated (Figure 2D). Thus, identified SLiMs in NUP153 and NOTCH1 mediate proximity, and likely direct interaction with CN *in vivo*.

**Figure 2:**
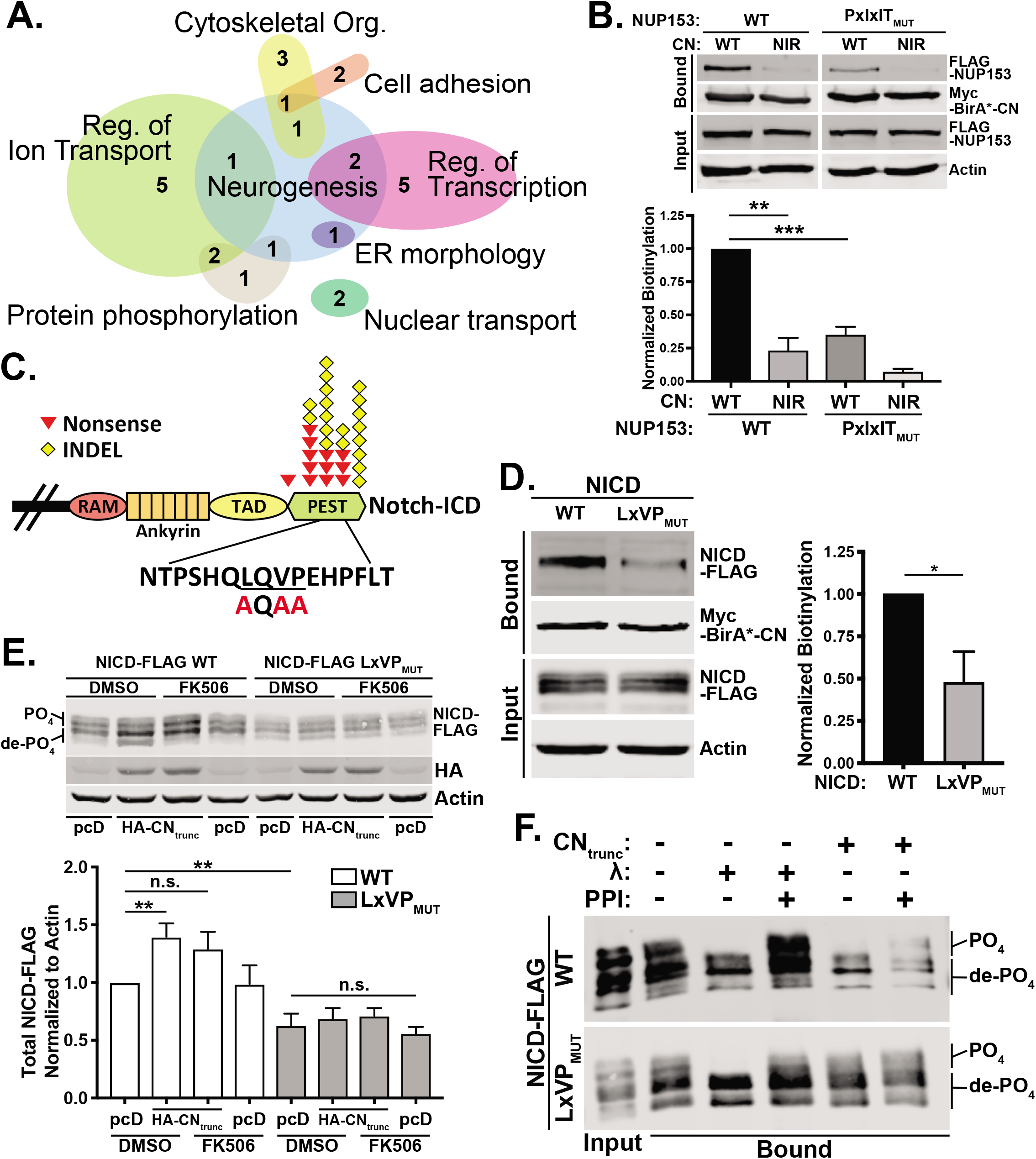
ProP-PD discovers CN interacting proteins and identifies NOTCH1 as a CN substrate. **A**. GO process terms associated with multiple CN SLiM-containing proteins identified via ProP-PD. **B**. NUP153 proximity to CN is PxIxIT-dependent. Biotinylated proteins were purified from HEK293T cells expressing myc-BirA*-CN (CN_WT_ or CN_NIR_) and FLAG-NUP153 WT (^485^**P**T**F**NF**S**^490^) or PxIxIT_MUT_ (^485^**<u>A</u>**T**<u>A</u>**NF**<u>A</u>**^490^) and probed with anti-FLAG antiserum. Data are mean ± SEM; n=3. **: p<0.01, ***: p<0.001. Blots were cropped to remove samples not relevant to these analyses. **C**. Schematic of NOTCH1 intracellular domain (NICD) showing sequence of WT and LxVP_MUT_. RAM: RBP-Jκ associated module, TAD: trans-activating domain, PEST: proline, glutamine, serine and threonine-rich domain. Triangles and diamonds: positions of stabilizing mutations found in T-ALL patients. **D.** NICD proximity to CN is LxVP-dependent. BioID as in **B** with myc-BirA*-CN and NICD-FLAG WT (^2502^**L**Q**VP**^2505^) or LxVP_MUT_ (^2502^**<u>A</u>**Q**<u>AA</u>**^2505^). Data are mean ± SEM; n=3; *: p<0.05. **E**. CN regulates NICD phosphorylation and steady state levels *in vivo.* Immunoblot analysis of NICD-FLAG (WT and LxVP_MUT_) expressed in HEK293 T-REx cells under basal (pcD), CN activating (HA-CN_trunc_) or inhibitory (FK506) conditions. Steady state levels of total NICD-FLAG are quantified. Data are mean ± SEM, n=6. **: p<0.01, n.s.: not significant. **F**. CN dephosphoryates NICD *in vitro.* NICD-FLAG WT (upper panel) or LxVP_MUT_ (lower panel) was purified from HEK293 T-REx cells and treated *in vitro* with CN_trunc_ or λ phosphatase in the presence or absence of phosphatase inhibitors (PPI), as indicated. See also Table S1.

To examine whether NOTCH1 is a CN substrate, NICD-FLAG was co-expressed with CN_trunc_, a constitutively active enzyme that lacks autoinhibitory sequences. This resulted in increased electrophoretic mobility of NICD_WT_ but not NICD-LxVP_MUT_, which was counteracted by the CN inhibitor, FK506 (Figure 2E). Incubating immunopurified NICD_WT_-FLAG with CNtrunc or the non-specific λ phosphatase *in vitro* also increased electrophoretic mobility via dephosphorylation, as it was blocked by phosphatase inhibitors (Figure 2F). In contrast, λ but not CNtrunc dephosphorylated NICD-LxVP_MUT_. Together, these data demonstrate SLiM-dependent dephosphorylation of NICD by CN *in vivo* and *in vitro* and establish NICD as a CN substrate. The C-terminal PEST domain of NICD, which is frequently mutated in T-cell Acute Lymphoblastic Leukemia, contains the LxVP motif and phosphorylation-dependent degradation signals (Figure 2C) (Tzoneva and Ferrando, 2012). We noted that steady-state levels of NICD-LxVP_MUT_ were lower than NICD_WT_. Also, levels of NICD_WT_, but not -LxVP_MUT_, increased with CNtrunc expression, although FK506 failed to reverse this effect (Figure 2E). Thus, CN may stabilize NICD – a possibility that aligns with findings that CN promotes Notch signaling *in vivo* (Kasahara et al., 2013; Mammucari et al., 2005; Medyouf et al., 2007). Importantly, these analyses support the use of SLiM discovery to elucidate CN signaling.

### Development of in silico methods for proteome-wide discovery of CN-binding SLiMs

Next, all validated CN-binding sequences were used to construct Position-Specific Scoring Matrices (PSSMs), statistical models that include weighted sequence information at each motif position and reflect the physiochemical properties of both core and flanking residues (Figure 3A-C; Table S2) (Li et al., 2007; Nguyen et al., 2019; Sheftic et al., 2016). Assessment of each PSSM by leave-one-out cross-validation showed that they were robust, sensitive, and specific (Figure S3A, B).

**Figure 3:**
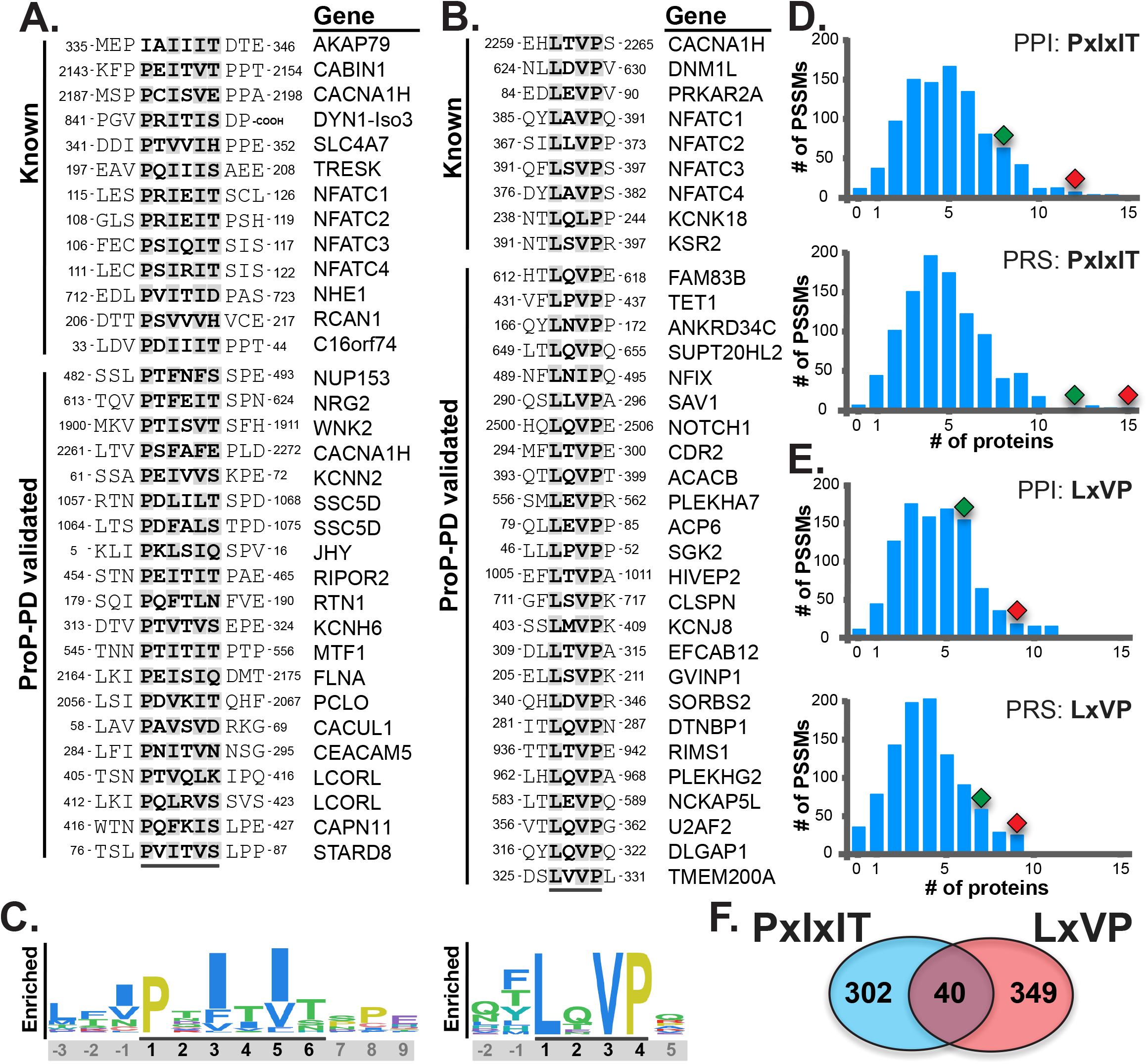
In silico identification of human CN-binding SLiMs. **A**, **B**. PxIxIT (A) and LxVP (B) sequences that were previously known or shown in this study. **C**. Schematic of PSSMs for PxIxIT and LxVP; residue size is proportional to its weighted value. **D**, **E**. PSSM validation. Blue bars: frequency distribution showing number of SLiM-containing proteins discovered by 1,000 randomly shuffled PSSMs in PPI (182 CN interacting proteins from public datasets) or PRS (104 CN substrates or interactors curated from literature). Red diamond: motif-containing proteins discovered by PxIxIT or LxVP PSSM. Green diamond: previously known motif-containing proteins. **F**. Number of human proteins containing CN SLiMs after filtering (see Fig. S3C). See also Figure S3 and Table S2.

To identify putative PxIxIT and LxVP motifs throughout the proteome, a series of filtering steps were applied (Figure S3C). All peptides matching a loose regular expression for each motif (P[^PG][FILV][^PG][FILV][^PG] or L.[ILV]P) were scored and ranked using the PSSM. Next, 1,121 PxIxIT and 1,167 LxVP instances with PSSM p-values ≤ 10^-4^ were filtered using additional criteria: 1) CN localizes to the cytosol and nucleus (Hallhuber et al., 2006), thus any extracellular, transmembrane, or lumenal sequences were removed; 2) SLiMs are found in regions of intrinsic disorder (Tompa et al., 2014) and > 90% of the training peptides have a high propensity for disorder. Therefore, sequences with IUPRED scores < 0.30 were removed (Dosztanyi et al., 2005), which significantly improved predictive power (Table S2). Conservation also modestly increased predictive power (Table S2), but was not used as a filter, as previous work showed rapid evolution of fungal PxIxIT motifs (Goldman et al., 2014).

The performance of each PSSM vs 1,000 randomly shuffled variants was measured on two sets of proteins that should be enriched for CN interactors: 1) The <u>P</u>rotein-<u>p</u>rotein <u>I</u>nteraction (PPI) dataset, which contains 182 CN interactors reported in public databases; 2) The <u>P</u>ositive <u>R</u>eference <u>S</u>et (PRS), 104 literature-curated proteins that are dephosphorylated by or interact with CN (Table S5). Both PxIxIT and LxVP PSSMs discovered a statistically enriched number of motif-containing proteins in each dataset and outperformed the majority of randomly shuffled PSSMs (Figure 3D, E; Table S2). Importantly, these analyses revealed motifs in known CN substrates, B-RAF, MEF2A, Amphiphysin and MAP3K7/TAK, some of which we confirmed to bind CN *in vitro* (Figure S3D). Taken together, these findings validate our *in silico* approach to CN target prediction.

Using this strategy, 409 putative PxIxITs and 410 putative LxVPs from 342 and 389 unique proteins, respectively, were identified in the human proteome, with only 40 proteins containing both motif types (Figure 3F). Because our strategy was optimized for selectivity rather than sensitivity, many additional proteins likely contain cryptic motifs that are functional, but not well scored by our PSSMs. In fact, multivalent SLiM-mediated interactions are often driven by a single high affinity/consensus motif in combination with degenerate variants that diverge significantly from the consensus (Ivarsson and Jemth, 2018; Stevers et al., 2018; Yaffe, 2002).

The prediction of hundreds of PxIxIT and LxVP instances in putative CN-interacting proteins across the human proteome suggests broad regulatory roles for CN, many of which are not represented in the current literature. Furthermore, because the SLiM-binding surfaces on CN are highly conserved, this *in silico* strategy can be generally applied, and we developed a web-based resource to identify putative CN-binding motifs in any protein or proteome of interest: http://slim.icr.ac.uk/motifs/calcineurin/.

### SLiM-dependent proximity labeling (BioID) identifies candidate CN interactors

Next, using proximity-dependent biotinylation coupled to mass spectrometry (PDB-MS)(Gingras et al., 2018), we explored the spatial distribution of CN within the cell, which should reflect the abundance and affinity of its SLiM-containing interacting partners. Biotinylated proteins were identified from HEK293 T-REx cells that stably express BirA*, BirA*-CN_WT_;, or docking mutants, -CN_NIR_ or -CN_WF_ (Figure 4A). BirA*-CN_WT_ specifically biotinylated 397 proteins (Table S3), including: known CN substrates and regulators (AKAP5, PRKAR2A, RCAN1 and NFAT4), interactors identified by ProP-PD (RTN1A and WNK2), as well as 50 proteins with SLiMs predicted *in silico* (Table S5). Importantly, proximity to CN was SLiM-dependent (Log_2_WT/Mut > 0.5) for almost half of these proteins (196/397), whose biotinylation by BirA*-CN_NIR_ (139) and/or -CN_WF_ (133) was reduced relative to BirA*-CN_WT_ (Figure 4B, S4A). PSSM analyses showed that PxIxIT and/or LxVP-containing proteins were statistically enriched in SLiM-dependent CN proximal proteins (Figure S4B, Table S2), but not in those that were SLiM-independent or in a set of proteins that are consistently labelled by BirA* alone (Table S2). Thus, BioID with WT vs. docking-defective BirA*-CN fusions combined with *in silico* identification of CN-binding SLiMs distinguishes candidate interactors from CN-proximal bystander proteins.

CN-proximal proteins were enriched for multiple cellular component GO Terms (Mi et al., 2013) that echo known regulation by CN of the cytoskeleton, vesicle trafficking and synaptic plasticity (Baumgartel and Mansuy, 2012; Cousin and Robinson, 2001; Hoffman et al., 2017)(Figure 4C, Table S3), and surprisingly identified multiple proteins at the centrosome and NPC – structures whose possible regulation by CN is unknown.

**Figure 4:**
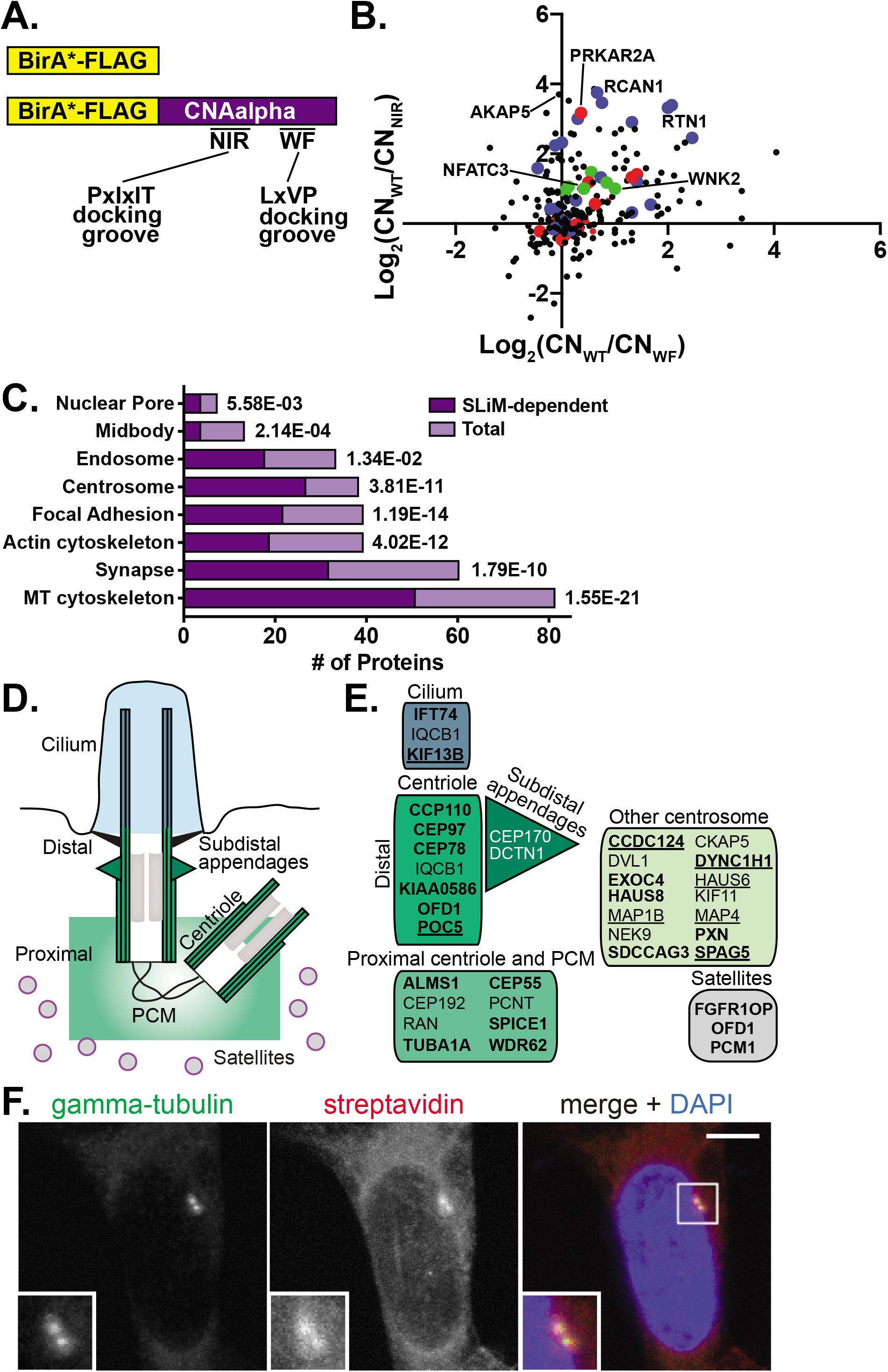
BioID identifies candidate CN-interacting proteins. **A.** Schematic of BirA* fusions to CNAa used for BioID. NIR and WF indicate CN residues required for PxIxIT and LxVP binding, respectively. **B**. Log_2_ ratio of spectral counts for proteins biotinylated by CN_WT_ or docking site mutants (CN_NIR_ and CN_WF_) and analyzed by data-dependent acquisition (DDA). Proteins containing PxIxIT (blue dots), LxVP (red dots) or both (green dots) are indicated. **C**. Select statistically enriched GO-cellular component categories shown for 397 CN-proximal (Total) or 196 ‘SLiM-dependent’ (Log_2_ WT/mutant > 0.5) proteins. **D**. Schematic of centrosome structure and associated regions including pericentrioloar material (PCM). **E**. Centrosomal location of CN-proximal proteins (Table S3). Bold: proteins with SLiM-dependent biotinylation. Underlined: proteins with a LxVP or PxIxIT identified by PSSMs (Table S4). **F**. Centrosomal distribution of biotin labeling in BirA*-CNAα -expressing HEK293 cells. Fluorescent microscopy shows γ-tubulin (green) at centrosomes, DAPI (blue) at nuclei and Streptavidin (red) marks sites of biotinylation. The region of interest is shown in inset. Images are a maximum projection of a confocal z-stack. Scale bar: 5 μm. See also Figure S4 and Table S3.

### CN shows proximity to centrosomal proteins

Centrosomes are the major microtubule-organizing centers in mammalian cells and contain two centrioles surrounded by pericentriolar material (PCM; Figure 4D)(Breslow and Holland, 2019). Centrioles are also required for cilia formation. We identified 35 centrosomal and/or ciliary proteins that were biotinylated by BirA*-CN_WT_, half of which (18) showed PxIxIT-dependence (Figure 4E, Table S3). In particular, proteins at the distal end of the centriole were highly represented, and were among those most strongly PxIxIT-dependent (CCP110, CEP97, CEP78, Talpid3/KIAA0586, OFD1 and POC5; Figure 4E, Table S3) (Azimzadeh et al., 2009; Singla et al., 2010; Tsang and Dynlacht, 2013). Furthermore, fluorescence microscopy showed biotin labeling at the centrosome in 99% of cells expressing BirA*-CN_WT_ (n=276), compared to 0% of cells expressing BirA* alone (n=185) or without added biotin (Figure S4C, D). This biotin signal co-localized with centrosomal proteins γ-tubulin (Figure 4F) and centrin in both interphase and mitotic cells (Figures S4C, D). Thus, these analyses revealed PxIxIT-dependent proximity of CN to centrosomal proteins that was not previously documented.

### CN co-localizes with and dephosphorylates NPC basket proteins

Bio-ID and ProP-PD analyses indicate possible functions for CN at the NPC, which is comprised of multiple copies of ~30 proteins (nucleoporins or nups) that serve structural roles and/or facilitate nucleocytoplasmic transport (Beck and Hurt, 2017). Nuclear transport proteins identified as CN-proximal and/or containing a CN-binding SLiM (Figure 5A) include soluble transport factors (Ran, RanGap1, RanBP1, Importin-β) and several nups, most of which directly participate in nuclear transport (Figure 5A). Consistent with our findings, while this paper was under review, multiple nups were detected as interacting with CN via AP-MS (Brauer et al., 2019).

**Figure 5:**
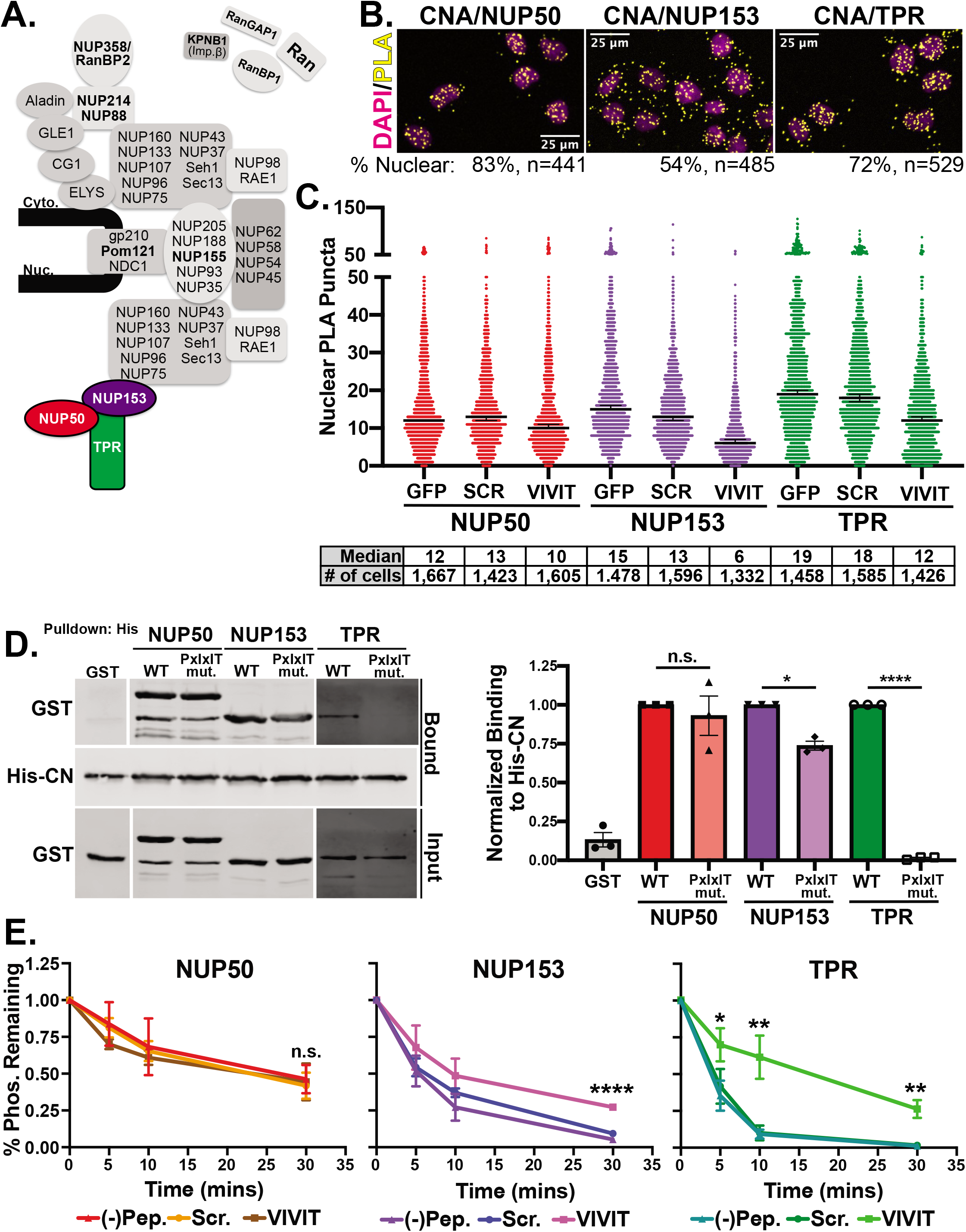
CN interacts with and dephosphorylates NPC basket proteins. **A**. Schematic of human NPC proteins, with nuclear basket nups in color. Bold: nups identified in PDB-MS and/or PSSM analyses. **B**. Proximity ligation assay (PLA) with endogenous CN and nuclear basket nups in HeLa cells. PLA signal: yellow, DAPI: magenta. Representative images from a single z-plane are shown and % PLA puncta overlaying the nucleus is indicated below. # of cells analyzed ranged from 421 to 529 across replicates (n=2) and conditions. Quantification in Figure S5A. **C**. PLAs performed as in B, with cells expressing GFP fused to VIVIT, SCR, or no peptide. Data are represented as median nuclear PLA puncta with 95% confidence intervals. Median values and # of cells analyzed are shown below each condition, n=3 replicates. Quantification and statistical analyses in Figure S5C and S5D. **D**. *In vitro* co-purification of GST-NUP50 (full length), -NUP153_228-611_ and -TPR_1626-2363_ WT or PxIxIT mutants with HIS-CN shows PxIxIT-dependence for CN-NUP153 and CN-TPR. Data are mean ± SEM; n=3, n.s.: not significant, *: p<0.05, ****: p<0.0001. **E.** ERK2-phosphorylated GST-NUP50, -NUP153_228-611_ and -TPR_1626-2363_ are de-phosphorylated by CN_trunc_ *in vitro*. Reactions contained peptides (VIVIT or SCR), as indicated. Phosphorylation (^32^P content) shown at indicated times relative to T=0. Data are mean ±SEM, n=3. n.s.: not significant, *: p<0.05, **: p<0.01. ****: p<0.0001. See also Figure S5.

In particular, we identified all three components of the NPC nuclear basket structure: NUP153 contains a validated PxIxIT (Figure 1G) and shows SLiM-dependent proximity to CN *in vivo* (Figure 2B), whereas NUP50 and TPR contain *in silico*-predicted PxIxIT sites (Table S2). To investigate possible association of CN with basket proteins *in situ,* we used Proximity Ligation Assays (PLA), which detect and amplify endogenous protein-protein interactions in fixed cells at <40 nm resolution (Gullberg et al., 2004). HeLa cells incubated with pairs of antibodies against CN and either NUP50, NUP153, or TPR, displayed robust PLA puncta, which substantially overlay the nucleus (Figure 5B and S5A). In contrast, cells incubated with either a single or no antibody showed very few puncta, as expected (Figure S5A). Next, we used PLA to measure CN-basket association in cells expressing GFP fused to either VIVIT, the high affinity PxIxIT peptide (Aramburu et al., 1999), SCR, a peptide in which the VIVIT sequence is scrambled, or GFP alone. These peptide fusions were distributed throughout the cytosol and nucleus (data not shown) and expressed at similar levels (Figure S5B). In each case, PLA puncta were significantly reduced by expression of VIVIT vs. SCR, with larger differences observed for CN-NUP153 and CN-TPR than for CN-NUP50 (Figure 5C, S5C and D). These results suggest that, *in vivo*, association of CN with the NPC basket is PxIxIT-dependent.

Next, we examined purified, recombinant basket nups for direct binding to CN. NUP153 and TPR contain functional PxIxIT sequences (Figure S5E) that, when mutated, reduce copurification of each protein with CN (Figure 5D). In contrast, the predicted PxIxIT in NUP50 failed to bind CN *in vitro* and mutating this site did not affect co-purification of full length NUP50 with CN (Figure 5D, S5E). Thus, although we observe CN binding to all 3 NPC basket proteins, the precise CN-binding motif(s) in NUP50 are yet to be identified.

Previous studies show that ERK phosphorylates multiple nups to inhibit nuclear transport (Kodiha et al., 2009; Kosako et al., 2009; Porter et al., 2010; Stuart et al., 2015); however, its opposing phosphatase is yet to be identified. Therefore, we investigated whether CN dephosphorylates ERK phosphosites *in vitro*. Using recombinant, ERK-phosphorylated NUP50, NUP153_228-611_, and TPR_1626-2363_, we observed robust dephosphorylation by CN_trunc_. Importantly, for NUP153 and TPR, but not NUP50, dephosphorylation was specifically inhibited by VIVIT (Figure 5E). Thus, NPC basket proteins are CN substrates and NUP153 and TPR, but not NUP50 contain functional PxIxITs. *In vivo*, effects of VIVIT on NUP50-CN association (Figure 5C) suggest that PxIxITs in NUP153, TPR and/or other NPC proteins may promote targeting of CN to NUP50 in the context of an assembled NPC.

### CN regulates NPC proteins and nuclear transport in vivo

To probe for CN effects on NPC function, we next examined nuclear accumulation of an NLS-containing reporter protein, RGG, in HeLa cells, in which ERK1/2 are active (data not shown). RGG contains GFP fused to an NLS that is activated by dexamethasone (DEX) (Love et al., 1998). After addition of DEX, live-cell imaging under control (DMSO) or inhibitory (FK506 or CsA) conditions showed that blocking CN activity decreased the rate and total amount of RGG accumulated in the nucleus (Figure 6A, B, Movie S1). Effects of FK506 and CsA, which inhibit CN through distinct mechanisms (Liu, 1993), were indistinguishable. Together, our *in vivo* and *in vitro* findings support the hypothesis that CN promotes nuclear accumulation of transport cargo by dephosphorylating NPC components.

**Figure 6:**
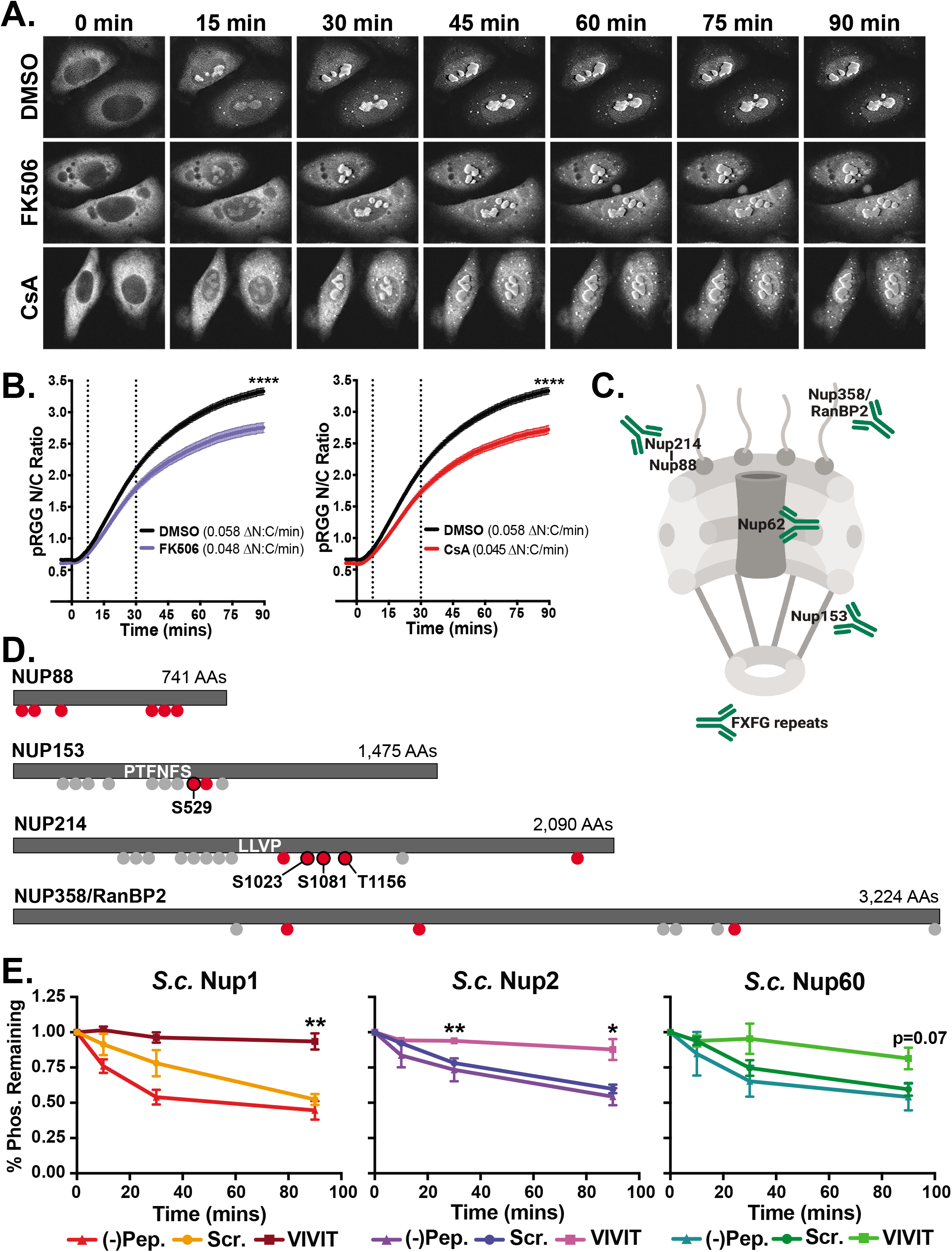
CN regulates nuclear transport and nup phosphorylation in vivo. **A, B.** Nuclear accumulation of RGG reporter in control (DMSO) or CN inhibited (FK506, CsA) HeLa cells. A. Frames from representative videos. B. Nuclear to cytoplasmic (N:C) ratio of GFP at indicated times after dexamethasone addition. Data are mean ± SEM for 146 (DMSO), 99 (FK506), or 81 (CsA) cells. Initial import rates, indicated as change in RGG N:C ratio over time (ΔN:C/min), were calculated from data within dashed lines. Extent of accumulation is significantly different between DMSO and FK506 or CsA (**** = p<0.0001). **C**. Nuclear pore schematic indicating FxFG repeat-containing proteins recognized by mAb414. **D**. Proteins purified from HeLa cells with mAb414 and analyzed by MS. Grey circles = CN-independent phosphosites; Red circles = sites whose phosphorylation was ≥ 1.2-fold increased for 2 out of 3 replicates in CsA vs. DMSO. Red circle, black outline = sites with increased phosphorylation in CsA (3/3 replicates, ≥ 1.2-fold increase) and p<0.057. **E**. Yeast nups are PxIxIT-dependent CN substrates: GST-NUP1, −2 and −60 phosphorylated by Hog1 and dephosphorylated with yeast CN_trunc_ in the absence of peptide (-Pep.) or with SCR or VIVIT. Phosphorylation was normalized to total protein at each time point. Data are mean ±SEM, n=3. *: p<0.05, **: p<0.01. See also Figures S6, S7 and Table S4.

To further investigate this, we analyzed the phosphorylation state of select nups under the same conditions as our nuclear transport assays. Using an antibody that recognizes FxFG repeat-containing nups (Davis and Blobel, 1987), which are required for nucleocytoplasmic transport, NUP153, NUp358/RanBP2 and the NUP214-NUP88 complex were purified from HeLa cells pre-treated with CsA or DMSO (Figure 6C). Each of these nups contained multiple phosphopeptides that were consistently enriched (≥ 1.2-fold) in CsA-treated cells (Figure 6D, S7A). Phosphosites that were significantly enhanced by CsA (p < 0.057) included known ERK sites (NUP153 S529 and NUP214 S1081 (Galan et al., 2014; Kosako et al., 2009)) as well as additional uncharacterized sites in NUP214 (S1023, T1156) (Figure 6D, S6, S7A, Table S4). In summary, our data show that CN binds and dephosphorylates nuclear basket nups *in vitro* and that nuclear transport and nup phosphorylation are altered in cells upon CN inhibition. Taken together, these findings warrant further investigation into the precise mechanisms by which CN regulates the NPC.

Finally, we asked whether CN regulation of NPC proteins might be conserved in *S. cerevisiae,* where Nups 1, 2 and 60, orthologs of Nup153 and Nup50 (Beck and Hurt, 2017), contain predicted PxIxITs and/or CN-dependent phosphorylation sites (Goldman et al., 2014). First, we tested yeast CN (GFP-Cna1) for interaction with GST-tagged nups in yeast extracts and found co-purification of CN with Nup1, but not 2 or 60 (Figure S7B). The Nup1-CN interaction was PxIxIT-dependent, as it was disrupted by VIVIT, but not SCR (Figure S7B). During osmotic stress, CN is active, and yeast basket proteins are regulated by the Hog1/p38 MAPK (Guiney et al., 2015; Regot et al., 2013). Therefore, we tested whether Hog1-phosphorylated basket proteins were substrates of CN *in vitro.* In each case, yeast Nups displayed PxIxIT-dependent dephosphorylation that was inhibited by VIVIT, but not SCR (Figure 6E). Together, these data suggest that Nups 1, 2 and 60 contain functional PxIxITs, although lower affinity motifs in Nup2 and 60 may explain their failure to co-purify with CN from extracts. Overall, these analyses show that dephosphorylation of NPC components by CN is evolutionarily conserved.

### Assembly and analysis of a SLiM-based CN interaction network

CN is ubiquitously expressed and regulates a range of tissue-specific processes that have not been systematically investigated. Therefore, we assembled a high-confidence interaction network comprised of 486 unique proteins with experimental and/or *in silico* evidence of CN association (Figure 7A, Table S5). This network includes: proteins that contain a newly validated CN-binding SLiM, established substrates and interactors from the PRS, proteins from the PPI or PDB-MS data sets that contain putative SLiMs, and proteins with one or more high-confidence CN-binding motifs identified *in silico* (i.e. PSSM score > lowest ranking experimentally validated motif, Table S5). These 486 proteins define a network termed the CNome, which is significantly enriched for PPIs (p-value < 1.0×10^-16^ (Szklarczyk et al., 2017)) and for a variety of GO terms, as expected given the ubiquitous distribution of CN and Ca^2+^-dependent signaling machinery in humans (Figure 7B). These include terms related to phosphorylation, Ca^2+^-dependent signaling, transcription, and the cytoskeleton – all of which expand on CN functions that are described in the published literature (Table S6). Extensive roles for CN in cardiac and synaptic signaling, where Ca^2+^ and CN-dependent regulation are prominent, are also indicated (see Discussion, Figure 7C, D). Enrichment of centrosomal and NPC proteins in the CN interaction network is consistent with the PDB-MS and functional analyses presented here. In total, this network demonstrates the power of systematic SLiM identification to discover sites of CN action and serves as a resource for the Ca^2+^ signaling community.

**Figure 7:**
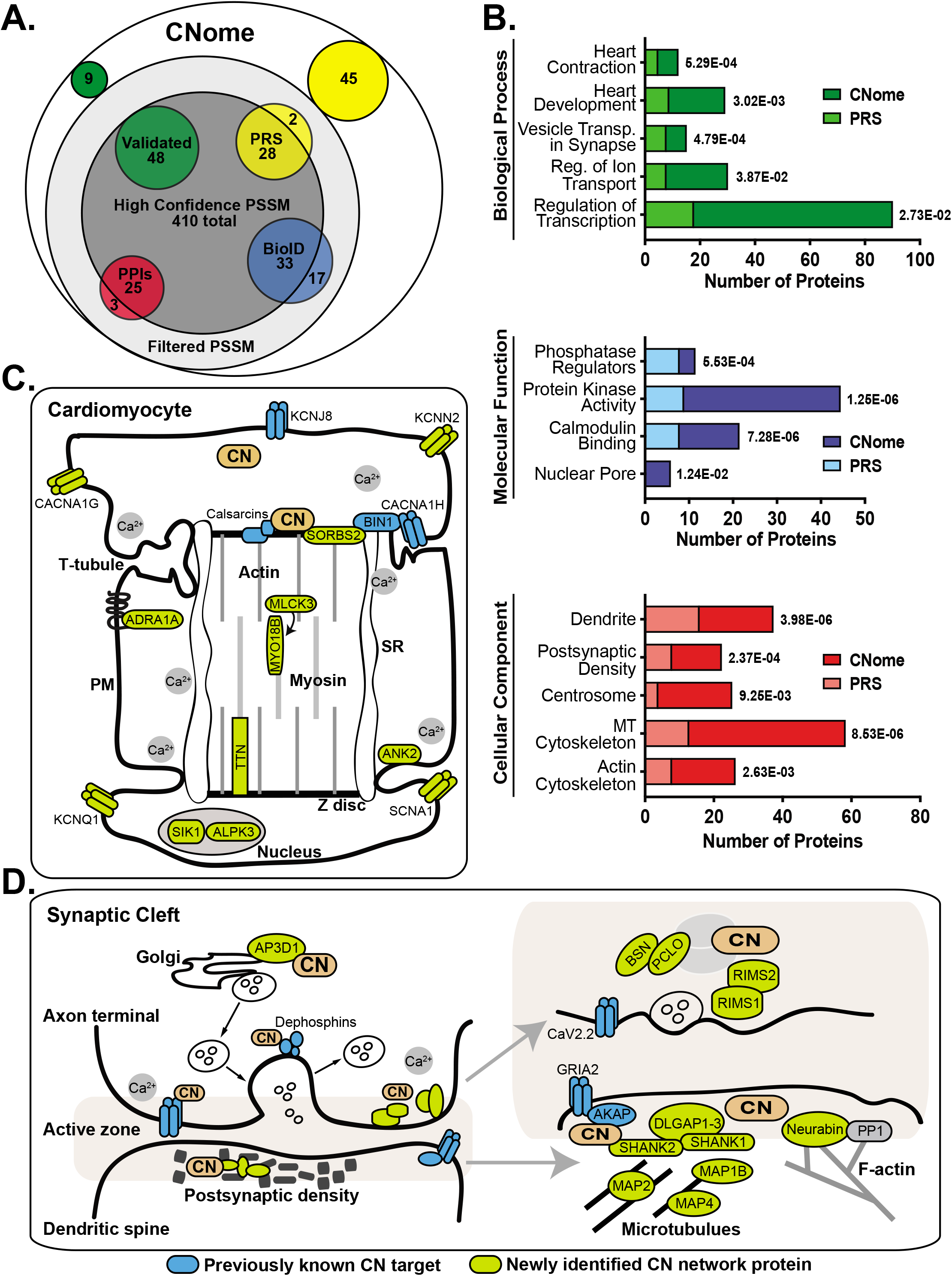
Functional analysis of SLiM-based CNome. **A**. 486 proteins of the CNome come from 6 datasets (see Table S4). Validated (green) = experimentally verified SLiMs (Figure 3A, B); PRS (yellow) = positive reference set; PPIs (orange) = high-throughput protein-protein interactions; BioID (blue)= CN-proximal proteins with SLiMs. Overlaps between Validated, PRS, PPI, and BioID datasets are not indicated in schematic (see Table S5). **B**. Significantly enriched GO terms in the CNome: Number of proteins for the CNome (dark color) vs. PRS (light color) are shown. FDR is indicated to the right of each bar. **C**, **D**. The CNome suggests novel targets for CN in cardiomyocytes (C) and neurons (D; presynaptic axons, the active zone and post synaptic dendrites). Gene names of established CN targets (blue) and novel CN network proteins (green) are shown. See Tables S5 and S6.

## Discussion

In this work, we focused on SLiM discovery and used both experimental and computational approaches to identify a CN network that provides surprising insights into signaling by this phosphatase.

Our approach to phosphatase substrate identification is fundamentally different from typical proteomic-based studies: Quantitative phosphoproteomics accurately identifies phosphorylation events that change when phosphatase activity is perturbed, but fails to distinguish direct from indirect effects, while AP-MS fails to capture many low-affinity, SLiM-mediated associations (Gingras et al., 2018). Furthermore, both approaches are limited by the choice of cell type or signaling conditions. In contrast, we leveraged ProP-PD to discover CN docking sites with amino acid resolution on a proteome-wide scale, proximity-based labeling (PDB-MS) to uncover transient CN docking interactions, and *in silico* specificity determinant modeling to pinpoint candidate interactors as well as key sequences that can be mutated to examine CN-mediated regulatory events across the proteome.

The PxIxIT and LxVP sequences described here identify unexpected CN-interacting proteins, including NOTCH1, NUP153 and TPR, each of which are directly dephosphorylated by CN *in vitro.* More importantly, this large collection of sequences was crucial for developing sensitive and robust statistical models (PSSMs) for systematic proteome-wide SLiM discovery. By encoding weighted information about each position of a motif, PSSMs offer significant advantages for motif identification over matching to a consensus sequence or regular expression (RegEX), which identifies only exact sequence matches and treats information at all motif positions equally (Krystkowiak et al., 2018). Limitations of this approach are illustrated by previous efforts to identify LxVP-containing proteins using a structure-based RegEx that fails to identify 16 of 25 experimentally validated LxVP-containing peptides discovered here via ProP-PD (Sheftic et al., 2016). Similarly, RegEx’s used for PxIxIT identification exclude information from flanking residues, which significantly influence binding affinity (Nguyen et al., 2019) and are represented in the 12 position PSSM developed here. A limitation of querying phosphatase function via SLiM discovery is the failure to identify a physiological context and/or tissue for candidate dephosphorylation events. However, these tools provide a powerful method to identify substrates in proteomic datasets, as shown here with PDB-MS studies.

The ability of PDB-MS to reveal subcellular organization is highlighted by the unexpected identification of SLiM-dependent CN proximity to proteins at the centrosome and especially at the distal end of centrioles, including CCP110 and CEP97. These proteins bind calmodulin and negatively regulate ciliogenesis (Spektor et al., 2007; Tsang et al., 2006). Previous reports of CN co-purification with Cep97 (Fogeron et al., 2013) and modulation of ciliary length by the CN regulator RCAN2 (Stevenson et al., 2018) suggest a role for CN at centrosomes/cilia. However, our as yet unsuccessful attempts to directly detect CN localization at centrosomes (data not shown) suggest that CN-centrosome interactions are limited, transient, and/or temporally regulated. Investigations of Ca^2+^ signaling at centrosomes, which contain multiple Ca^2+^- and/or calmodulin-binding proteins (Galletta et al., 2014; Khouj et al., 2019; Spektor et al., 2007; Tsang et al., 2006), and cilia, which constitute a distinct Ca^2+^ compartment (Delling et al., 2013; Delling et al., 2016), are ongoing. Our findings point to CN as a likely downstream effector of Ca^2+^ signals at these organelles.

The SLiM-based approaches used here also yielded unanticipated evidence that CN regulates nucleocytoplasmic transport and dephosphorylates NPC proteins. We showed that CN positively regulates the rate and accumulation of one transport cargo, although further studies are required to determine how broadly CN affects nucleocytoplasmic transport. Furthermore, because CN activity is strictly controlled by the availability of Ca^2+^ and calmodulin (Mehta and Zhang, 2014), our findings suggest that these signaling molecules are locally regulated at NPCs. The existence of calmodulin-dependent transport pathways provides evidence of functional calmodulin pools at NPCs (Wagstaff and Jans, 2009). Similarly, NPCs may represent a discrete Ca^2+^ signaling domain, as the nuclear envelope stores and releases Ca^2+^ with dynamics that are distinct from cytosolic Ca^2+^ signals (Bengtson and Bading, 2012; Ljubojevic and Bers, 2015). While previous work showed that changes in [Ca^2+^]_cyto_ alter the structure of key transport nucleoporins (Mooren et al., 2004; Paulillo et al., 2006; Sakiyama et al., 2017; Stoffler et al., 1999), results concerning the possible Ca^2+^-dependence of nucleocytoplasmic transport are conflicting (Greber and Gerace, 1995; Strubing and Clapham, 1999). Our findings argue that Ca^2+^ dynamics at NPCs should be re-examined. More broadly, our studies uncovered many potential CN substrates with functions that are currently unconnected to Ca^2+^ signaling, therefore providing rich new avenues for exploration of Ca^2+^-regulated processes.

Collectively, our analyses yield a catalogue of candidate targets of CN-dependent signaling throughout the body. This is exemplified by processes in the excitable tissues, heart and brain, where roles for Ca^2+^ and CN signaling are well-documented. CN regulates heart development and function via NFATs and signals from cardiac Z-discs, whose components are enriched in the CNome (Figure 7C) (Parra and Rothermel, 2017) (Chang et al., 2004; Wilkins and Molkentin, 2004). Moreover, CNome proteins align with the pleiotropic consequences of CN deletion in mouse cardiomyocytes (Maillet et al., 2010). Key signaling, structural and contractile proteins that contain CN-binding SLiMs may underlie defects in cardiac development, proliferation and contractility displayed by CN KO mice, while the enrichment of ion channels may explain observed arrhythmias. Similarly, CN is highly expressed in the brain, where it impacts learning, memory and other aspects of behavior (Baumgartel and Mansuy, 2012; Lin et al., 2003; Suh et al., 2013). The CNome suggests CN targets both pre- and post-synaptically, and particularly in the pre-synaptic active zone which coordinates neurotransmitter release (Figure 7D). These include BSN, PCLO, RIMS1, and RIMS2, which contain predicted SLiMS and show Ca^2+^-dependent phosphorylation decreases during exocytosis (Kohansal-Nodehi et al., 2016). In sum, the network presented here provides a road map for targeted elucidation of CN signaling pathways in the heart, brain and other tissues.

More evidence of undocumented CN signaling comes from patients chronically exposed to CN inhibitors (CNIs), which have been used as immunosuppressants for >35 years. The

CNome provides possible insights into adverse effects of CNIs, which include hypertension, diabetes, and seizures, and are caused by CN inhibition in non-immune tissues (Roy and Cyert, 2019). For example, CNI-induced hypertension results in part from hyperphosphorylation of the kidney Na/Cl cotransporter SLC12A3 (Hoorn et al., 2011), whose kinases (ASK3/MAP3K15, STK39/SPAK, and OXSR1/OSR; (Naguro et al., 2012)) have CN-binding SLiMs, as does NEDD4L, a regulator of another hypertension target, EnaC (Rizzo and Staub, 2015). Thus, the CNome may yield strategies to alleviate complications incurred during CNI-based immunosuppression. Similarly, this network contains many epilepsy-associated genes (Wang et al., 2017), and suggests mechanisms by which mutation of CNAa may cause this disease (Mizuguchi et al., 2018; Myers et al., 2017; Wang et al., 2017).

In summary, the CN signaling network (CNome) established here provides a basis for future investigations of Ca^2+^/CN signaling in healthy and diseased cells, including effects of prolonged immunosuppression. Moreover, our studies emphasize the importance of continuing to identify and characterize SLiMs, components of the dark proteome (Bitard-Feildel et al., 2018) that can unlock the mysteries of cell signaling. All major signaling pathways rely on SLiMs to form rapid, specific, and dynamic PPIs that transmit cellular information, and our studies provide a blueprint for investigating any SLiM-based network using a combination of experimental and computational strategies.

## Supporting information

Supplemental Table 1: ProP-PD Analyses

Supplemental Table 2: PSSM Benchmarking Table

Supplemental Table 3: BioID/MS Analyses

Supplemental Table 4: Nucleoporin Phosphoproteomics Analyses

Supplemental Table 5: CNome Composition and Assembly

Supplemental Table 6: GO Terms for the CNome

## Acknowledgements

We are indebted to Idil Ulengin-Talkish for critical feedback and thank Mark Dell’Acqua and Jeff Molkentin for helpful discussion, Jeremy Thorner and Francesc Posas for reagents, and the Skotheim lab for use of their cell culture facilities. We gratefully acknowledge NIH grants that funded MSC, JR, and NPD (R01 GM119336); DB (T32 GM007276); S-LX and MAK (R01 GM135706); CPW (F32 GM120916); KU and DM (1R01 GM131052); TS (R35 GM130286) and JTW (K99 GM131024). The Carnegie endowment funded S-LX, MAK, and SHH. ET was supported by a Stanford Graduate Fellowship. NED and IK were funded by Science Foundation Ireland Starting Investigator Research Grant 13/SIRG/2193; YI, VY and CB were funded by the Swedish research council (2016-04965), the Carl Trygger Foundation (CTS14:209) and Lenannders Foundation. ER acknowledges support from the Landesoffensive zur Entwicklung wissenschaftlich-ökonomischer Exzellenz (LOEWE), LOEWE-Zentrum Translationale Medizin und Pharmakologie. ACG and CJW were funded by the Canadian Institutes of Health Research (Foundation grant FDN143301) and the Governments of Canada and Ontario through Genome Canada, Ontario Genomics and the Ontario Research Fund (OGI-139 and RE08-065). ACG is the Canada Research Chair in Functional Proteomics.

## Author contributions

CPW and JR performed the majority of wet-lab experiments, supervised by MSC, with the exception of the following: VY and CB carried out the proteomic phage display screening, supervised by YI; ER performed the NGS analysis; CJW carried out BioID labeling, MS analyses and data processing, supervised by ACG; DM carried out live-cell imaging and data analyses of RGG accumulation, supervised by KU; JTW carried out imaging of biotin-labelled cells, and annotation of centrosomal proteins identified by BioID, supervised by TS. DB and ET analyzed PLA data. Bioinformatic analyses were carried out by NED, NPD and IK. S-LX, SHH and MK performed MS analyses, CPW, JR, MSC and NED conceived and supervised the work and wrote the manuscript with input from all the authors.

## Supplemental Items

**Figure S1: (related to Figure 1).**
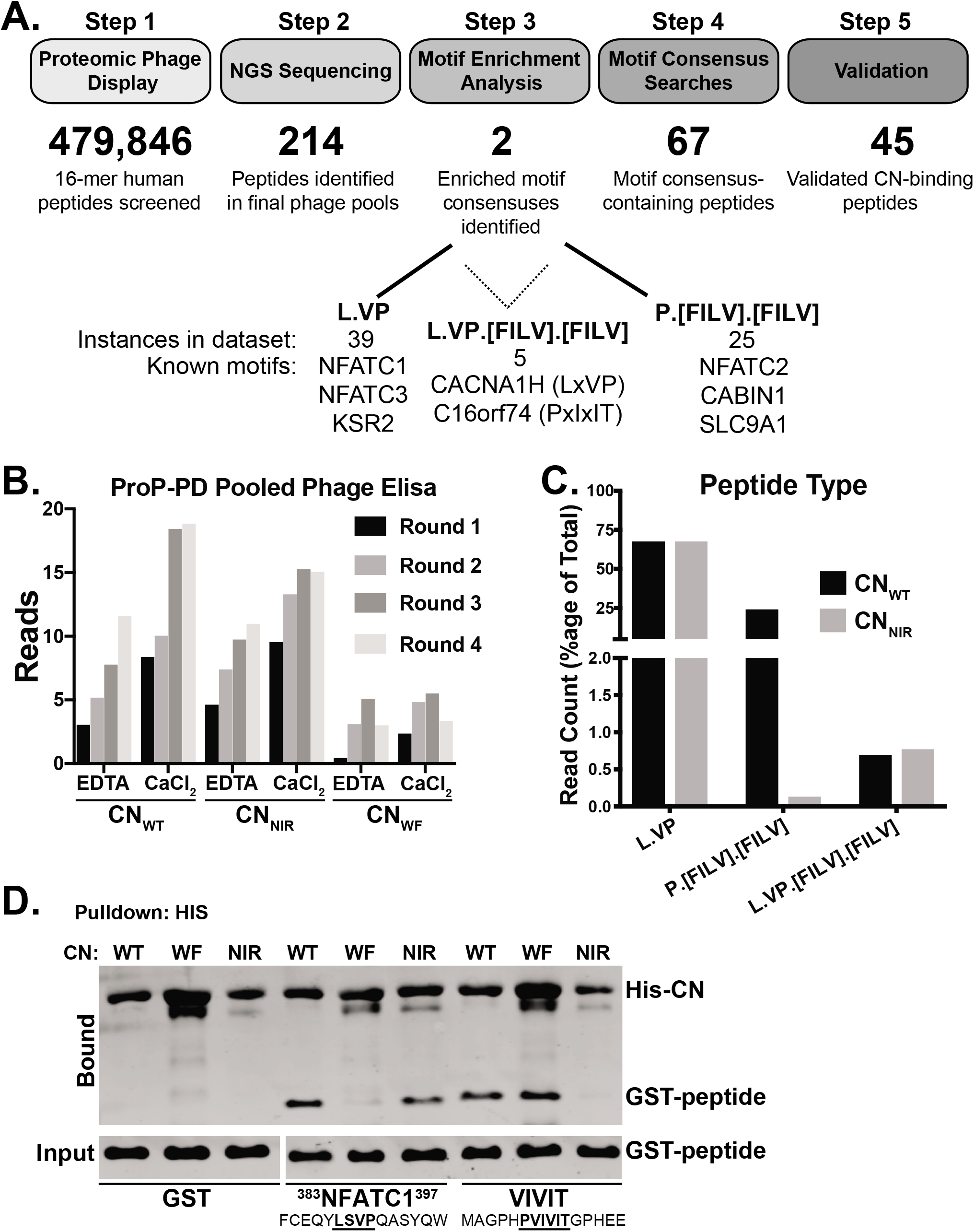
**A**. Flowchart for ProP-PD screen, motif enrichment and experimental validation of CN-Binding peptides. **B**. CN Sequence read counts for selected peptide-phage using WT or mutant calcineurin over 4 selection rounds are shown. Experiments were carried out in buffer containing CaCl_2_ or EDTA as indicated. CN_NIR_ contains ^330^NIR^332^-AAA mutations; CN_WF_ contains ^352^W-A, ^356^F-A mutations in CNAα. Phage selections were robust for CN_WT_ and CN_NIR_ but not for CN_WF_. **C**. Distribution of read counts for the indicated peptide type between CN_WT_ and CN_NIR_. **D**. *In vitro* binding of purified His-tagged CN_WT_, CN_NIR_ and CN_WF_ to GST-tagged LxVP peptide from NFATC1 or a high affinity PxIxIT peptide, VIVIT (Aramburu et al., 1999). CN_NIR_ has substantially reduced binding to VIVIT but not to NFATC1-LxVP whereas CN_WF_ has reduced binding to NFATC1-LXVP but not to VIVIT. See also Table S1.

**Figure S2: (related to Figure 1).**
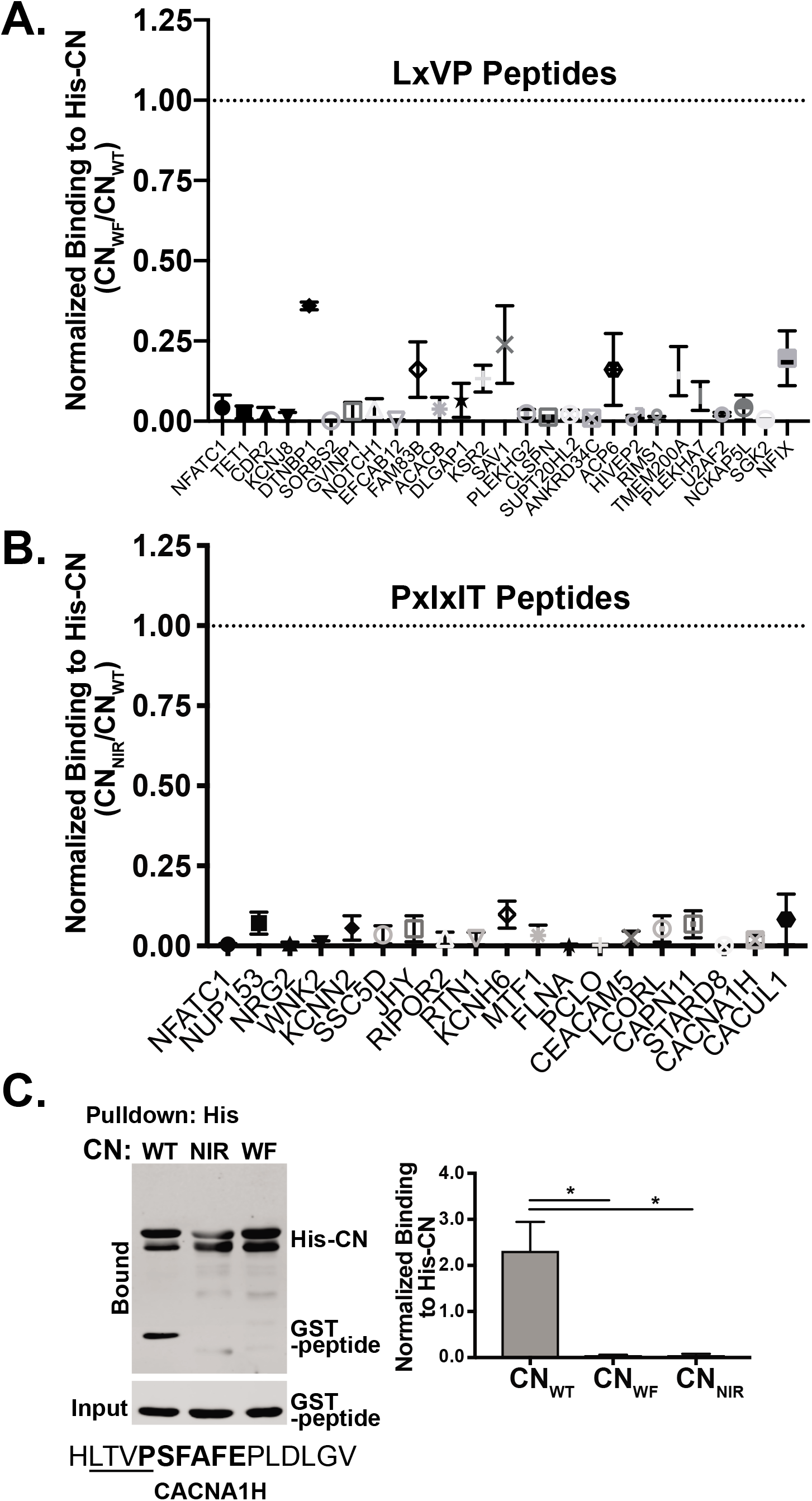
**A**. LxVP-peptides show greatly reduced binding to CN_WF_. *In vitro* co-purification of LxVP peptides with purified His-CN_WT_ and His-CN_WF_. Graph represents mean with SEM of two experiments. Binding to CN_WF_ is represented as a fraction of binding to CN_WT_, as indicated by dotted line. **B**. PxlxIT-peptides show greatly reduced binding to CN_NIR_. *In vitro* co-purification of PxIxIT peptides with purified His-CNwτ and His-CN_NIR_. Graph represents mean with SEM of two experiments. Binding to CN_NIR_ is represented as a fraction of binding to CN_WT_, as indicated by dotted line. **C**. *In vitro* co-purification of GST-tagged LxVPxlxIT peptide from CACNA1H with purified WT (CNWT) and mutant (CN_NIR_ or CN_WF_) His-CN heterodimer. Data are mean ± SEM, n=3, *: p<0.05. See also Table S1.

**Figure S3: (related to Figure 3).**
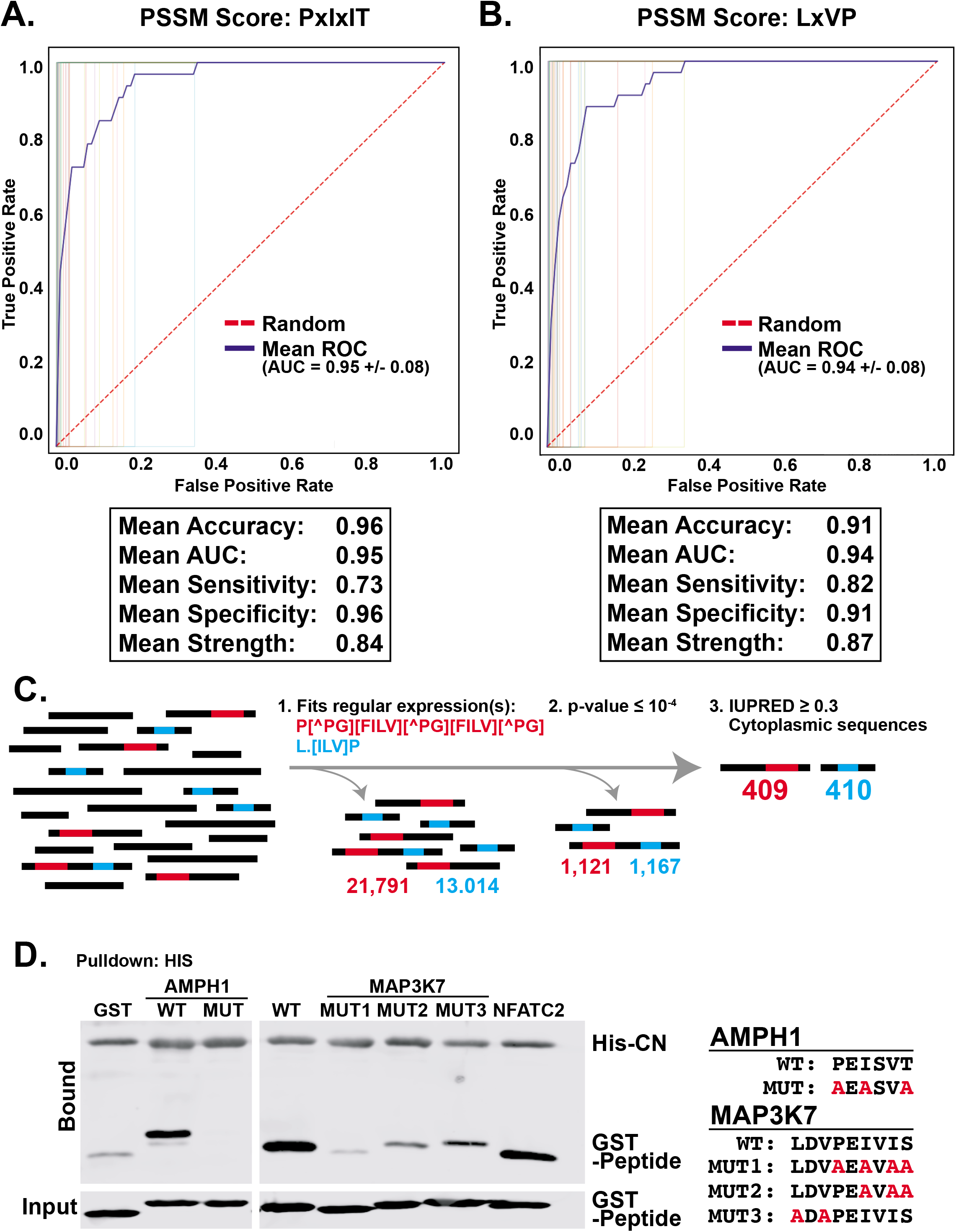
**A, B.** Receiver operating characteristic (ROC) curves for leave-one-out cross-validation of PxIxIT (A) and LxVP (B) PSSMs. The blue line represents mean PSSM values whereas the red line represents those for a random sequence. Benchmarking parameters for each PSSM are depicted below the curves. **C.** Overview of computational prediction of CN binding SLiMs from the human proteome. **D**. Newly predicted CN SLiMs in previously known CN regulated proteins (AMPH and MAP33K7) bind CN *in vitro.* Co-purification of GST-tagged WT CN SLiMs or mutant versions, as indicated, with purified His-CN_WT_. The PxIxIT SLiM (PEISVT) from AMPH, but not a mutant version, co-purifies with CN. M3K7 contains a combination LxVPxlxIT motif, where mutations in both the LxV and IxIT segments reduce interactions with CN_WT_. See also Table S2.

**Figure S4: (related to Figure 4).**
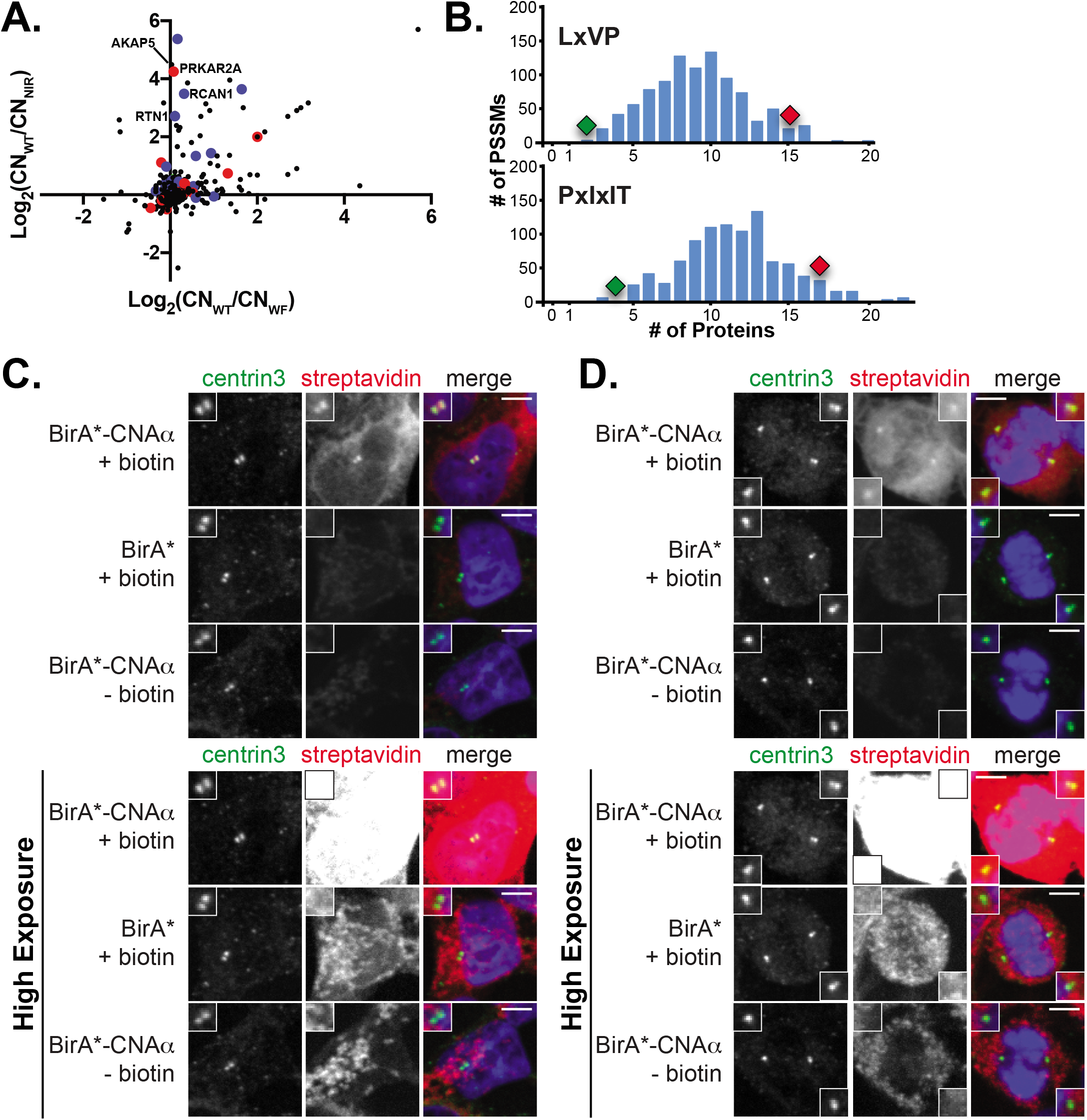
**A**. Log_2_ ratio of spectral counts of proteins biotinylated by CN_WT_ or docking site mutants (CN_NIR_ and CN_WF_) in HEK293 Flp-In T-REx cells. Peptides were analyzed by sequential window acquisition of theoretical fragments (SWATH, data-independent acquisition). Proteins containing PxIxIT (blue dots), LxVP (red dots) or both (green dots) SLiMs are indicated. **B.** Frequency of SLiM-containing proteins discovered for 1,000 randomly shuffled PSSMs in the set of 196 proteins that show CN docking-site dependent biotinylation. Red diamond: number of proteins with newly identified CN SLiMs. Green diamond: number of proteins with previously known CN SLiMs. **C, D.**: BirA*-CNA_WT_ signal co-localizes with centrosomes. HEK-293 Flp-In T-REx cells either at interphase (C) or undergoing mitosis (D), expressing BirA*-CNA_WT_ were incubated with biotin. Negative controls were processed in parallel: biotin-treated cells expressing BirA* alone, and untreated cells expressing BirA*-CNA_WT_. Centrosomes are marked by centrin3 (green). Streptavidin (red) binds to proteins labeled by BirA*-CNAWT or BirA* alone. DAPI (blue) marks the nucleus. Images are maximum projections of confocal z-stacks. Scale bars: 5 μm. Lower panels are the same images with enhanced streptavidin signal. See also Table S3.

**Figure S5: (related to Figure 5).**
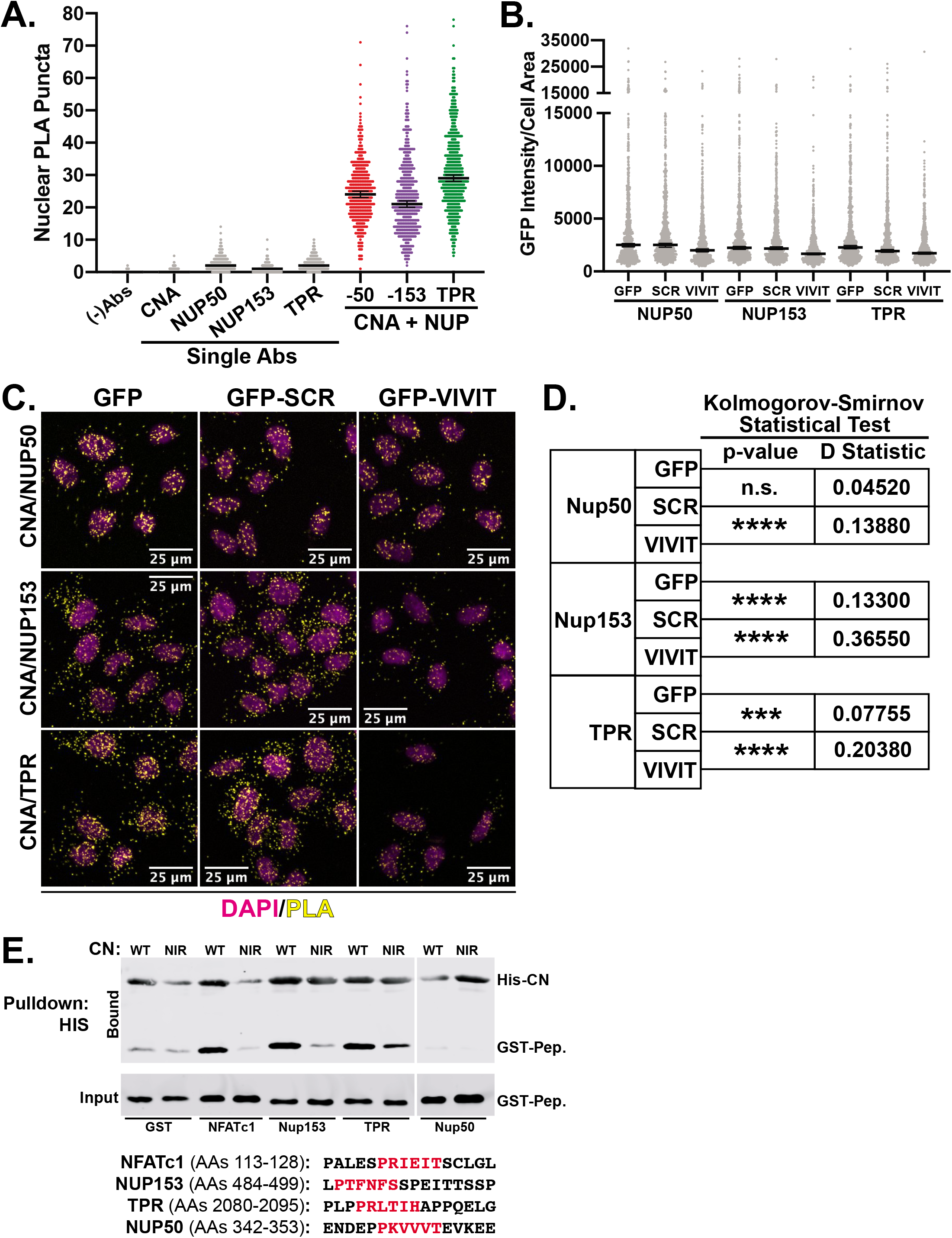
**A**. Proximity ligation assays (PLAs) with CNA and nuclear basket nup antibodies in HeLa cells. Median nuclear PLA puncta is indicated with 95% confidence intervals, n=2 biological replicates. **B**. Expression levels of GFP-peptide fusions in HeLa cells depicted in C and quantified in Figure 5C. **C.** Representative images of PLAs with CNA and nuclear basket nups in HeLa cells expressing GFP or GFP-peptides (SCR or VIVIT). Yellow: PLA signal; Magenta: DAPI. Images are from a single z-plane. **D**. Statistical analyses of cells from C, showing significantly decreased nuclear PLA puncta cells expressing GFP-VIVIT vs -SCR. **E.** Nuclear basket proteins contain CN-binding SLiMs. Co-purification of GST-tagged PxIxIT peptides with purified His-CN_WT_ or His-CN_NIR_. NUP153 and TPR PxIxIT peptides copurify with CN *in vitro*. Co-purification is PxIxIT-dependent and diminished with CN_NIR_.

**Figure S6: (related to Figure 6).**
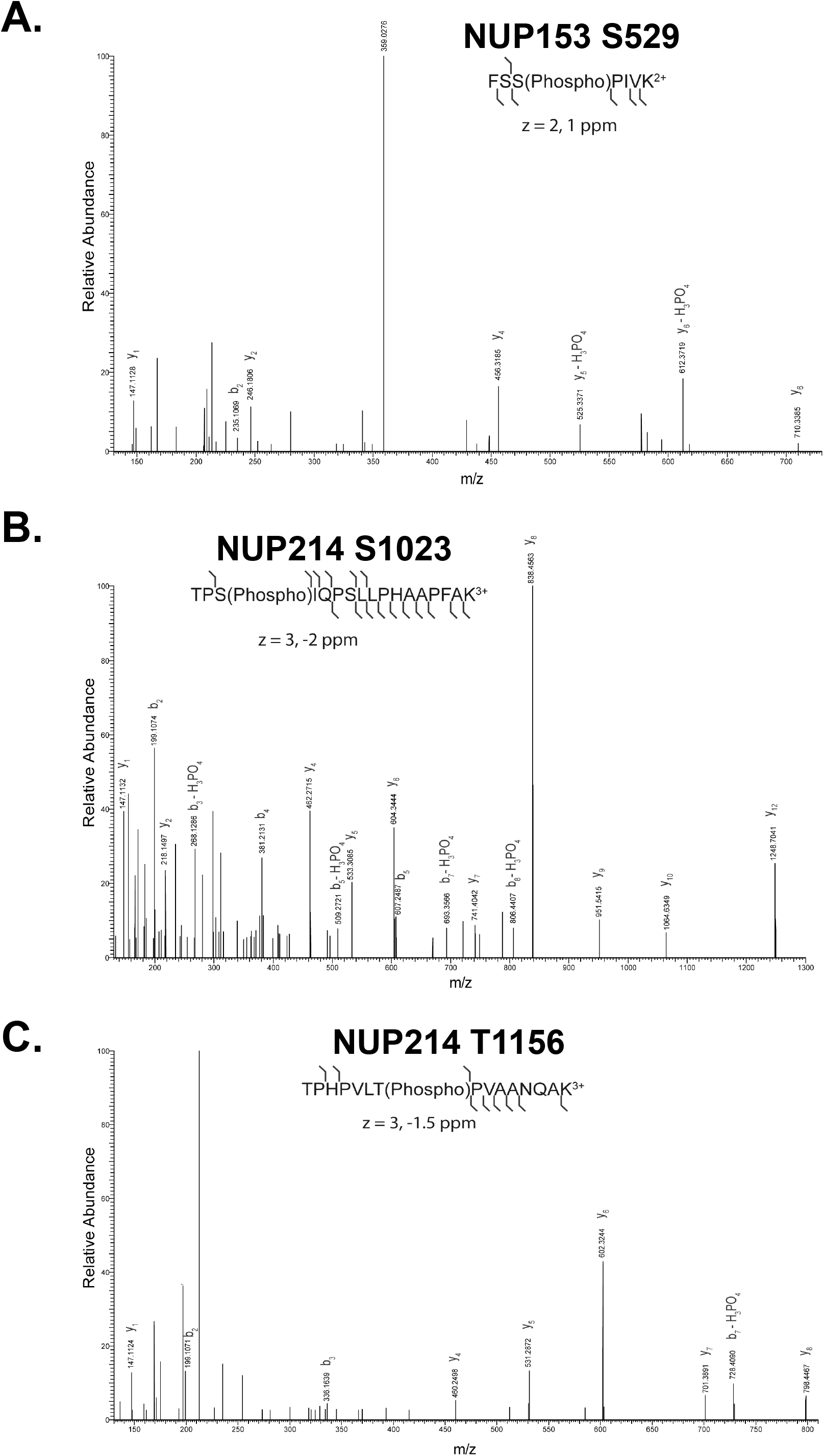
CN regulates NUP phosphorylation sites *in vivo.* Higher energy collision dissociation (HCD) mass spectra showing phosphorylation of serine 529 in Nup153 (**A**), serine 1023 (**B**) and threonine 1156 (**C**) in Nup214. See also Table S4.

**Figure S7: (related to Figure 6).**
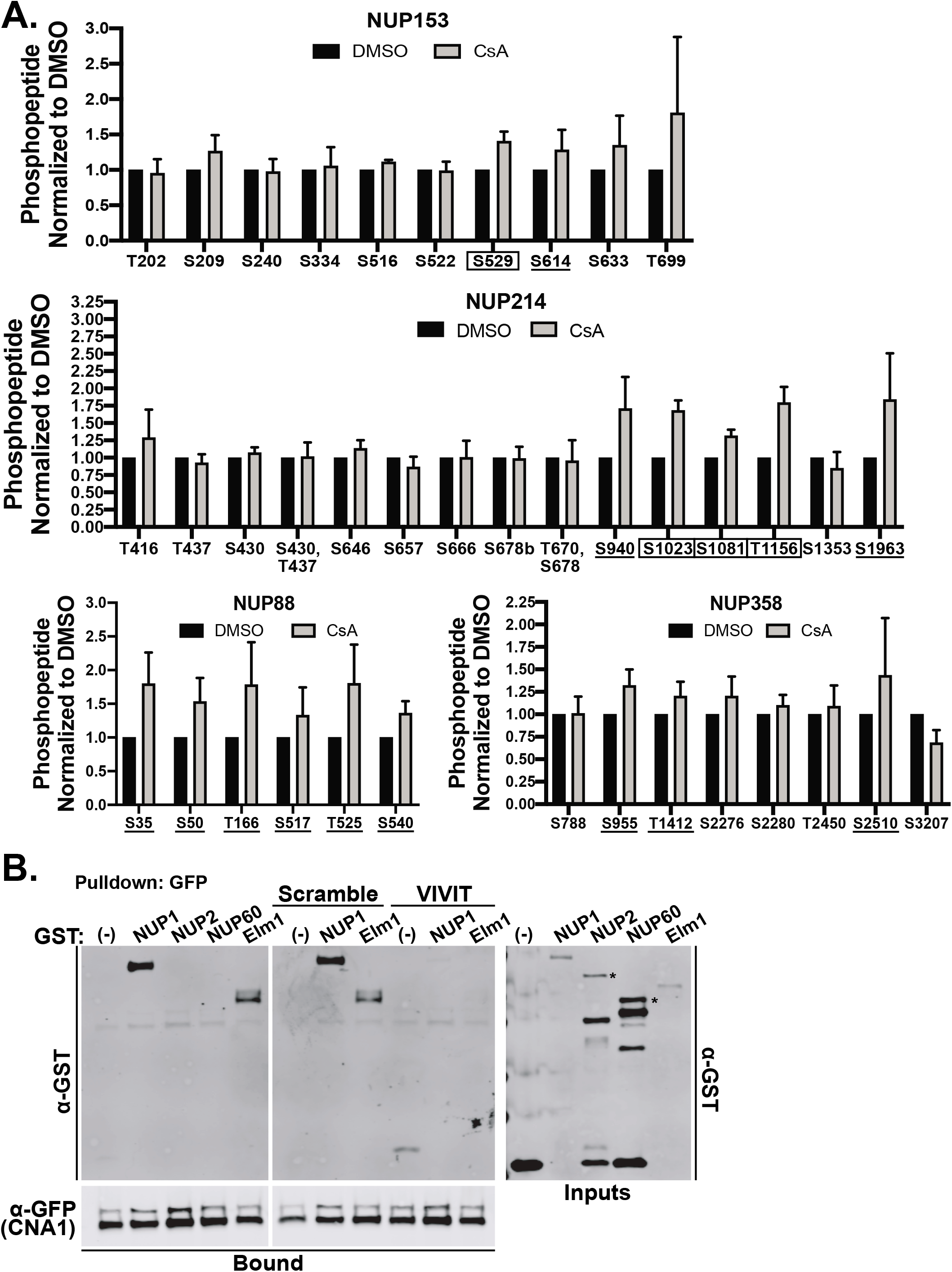
**A.** Quantification of nucleoporin phosphosites under CN inhibited conditions (CsA) normalized to control (DMSO). Underlined: sites where CN inhibition increased phosphorylation by at least 1.2-fold in 2 out of 3 replicates. Boxed: Sites where increase in phosphorylation was significant in 3 out of 3 replicates, p<0.057. **B.** Yeast CN interaction with Nup1 is PxIxIT-dependent. Left panel: Co-purification of GST-Nup1 with GFP-CNA1 from yeast extracts (JRY19) expressing indicated GST fusion proteins. Elm1 is a known CN interactor. Middle panel: Addition of VIVIT, a high-affinity PxIxIT peptide vs Scramble (negative control) disrupts PxIxIT-dependent CN binding to NUP1 and Elm1. Inputs: * indicates full-length proteins. See also Table S4.

## Additional information is presented in supplemental tables (S1-S5) and movie S1

**Table S1:** *ProP-PD results and analyses* (related to Figure 1). ProP-PD read counts, motif enrichment analyses, and in vitro binding assay results. Previously known CN-binding SLiMs are highlighted in yellow.

**Table S2:** *PSSM results and benchmarking* (related to Figure 3). Final filtered and unfiltered PSSM datasets as well as PSSM assembly, benchmarking, and interaction datasets.

**Table S3:** *BioID/MS analyses and GO Terms* (related to Figure 4). BioID/MS data-dependent (DDA) and -independent (SWATH) datasets and associated GO Term analyses. MassIVE SWATH 1: sample description table. MassIVE SWATH 2: protein identification evidence. MassIVE SWATH 3: MassIVE SAINTexpress v.3.6.1 output. MassIVE DDA 1: sample description table. MassIVE DDA 2: protein identification evidence. MassIVE DDA 3: MassIVE SAINTexpress v.3.6.1 output.

**Table S4:** *Phosphoproteomic analyses on immunopurified NPC proteins* (related to Figure 6). Raw and normalized results from skyline analyses performed on immunopurified nups from HeLa cells treated with DMSO or CsA.

**Movie S1:** *CN regulates accumulation of a nuclear transport reporter* (related to Figure 6) Nuclear accumulation of the dexamethasone-responsive RGG reporter was imaged in HeLa cells pre-treated with DMSO or CN inhibitors, FK506 or CsA (1 μM). Images of GFP signal were collected every 30 seconds from −5 minutes to +89.5 minutes after addition of dexamethasone). Two representative cells are shown from each condition. Quantitative analyses (Figure 6B) were performed with n = 146 (DMSO), 99 (FK506), or 81 (CsA) individual cells.

**Table S5:** *CNome assembly and composition* (related to Figure 7). Individual datasets used to assemble the final CNome. The final assignments and lists used for GO Term analyses are on the last tab.

**Table S6:** *CNome GO Terms* (related to Figure 7). GO Term analyses performed on the CNome. Proteins also found in the PosRefSet are highlighted in yellow.

## STAR Methods

### Purification of Calcineurin

For proteomic peptide phage display, GST tagged human calcineurin A (α isoform, truncated at residue 400), either wild-type or mutants ^330^NIR^332^-AAA and ^352^W-A,^356^F-A were purified. Constitutively active, truncated calcineurin was used to ensure that both PxIXIT and LxVP surfaces were accessible. Calcineurin A subunits were expressed in tandem with the calcineurin B subunit in *E. coli* BL21 (DE3) cells (Invitrogen) and cultures in LB medium containing carbenicillin (50 μg/ml) at 37°C to mid-log phase. Expression was induced with 1 mM isopropyl 1-thio-β-D-galactosidase (IPTG) at 16°C for 18 hours. Cells were pelleted, washed and frozen at −80°C for at least 12 hours. Thawed cell pellets were re-suspended in lysis buffer (50 mM Tris-HCl pH 8, 2 mM EDTA, 2mM EGTA, 150 mM NaCl, 1mM dithiothreitol (DTT), protease inhibitors) and lysed by sonication using four, 1-minute pulses at 40% output. NaCl added to 1.50M and extracts were clarified using two rounds of centrifugation (20,000 X g, 20 min). Tween-20 was added to 0.1% and extracts were bound to 1 ml of glutathione-sepharose beads (Fisher Scientific, USA) for 2-4 hr. at 4°C, in batch. Bound beads were loaded onto a column and washed with 10 column volumes of wash buffer (100 mM Tris-HCl pH 8.0, 100 mM KOAc, 2 mM MgOAc, 0.005% Tween-20, 1 mM DTT) followed by one column volume of wash buffer with 1 mM ATP and then eluted in wash buffer containing 40 mM glutathione, pH 8.0 and 5mM DTT. Purified calcineurin heterodimer were dialysed in buffer (50 mM Tris-HCl pH 7.5, 150 mM NaCl, 1 mM DTT) and stored in 10-15% glycerol at −80°C.

For *in vitro* peptide binding assays, N-terminally, 6-His-tagged human calcineurin A (α isoform, truncated at residue 400), either wild-type or mutants ^330^NIR^332^-AAA and ^352^W-A, ^356^F-A were expressed in tandem with the calcineurin B subunit in *E. coli* BL21 (DE3) cells (Invitrogen, USA) and cultured in LB medium containing carbenicillin (50 μg/ml) at 37°C to midlog phase. Expression was induced with 1 mM IPTG at 16°C for 18 hours. Cells were pelleted, washed and frozen at −80°C for at least 12 hours. Thawed cell pellets were re-suspended in lysis buffer (50 mM Tris-HCl pH 7.5, 150 mM NaCl, 0.1% Tween 20, 1mM β-mercaptoethanol, protease inhibitors) and lysed by sonication using four, 1-minute pulses at 40% output. Extracts were clarified using two rounds of centrifugation (20,000 X g, 20 min) and then bound to 1 ml of Ni-NTA agarose beads (Invitrogen) in lysis buffer containing 5mM imidazole for 2-4 hr. at 4°C, in batch. Bound beads were loaded onto a column and washed with lysis buffer containing 20mM imidazole and eluted with lysis buffer containing 300 mM imidazole, pH 7.5. Purified calcineurin heterodimer were dialyzed in buffer (50 mM Tris-HCl pH 7.5, 150 mM NaCl, 1 mM β-mercaptoethanol) and stored in 10-15% glycerol at −80°C.

### Proteomic peptide phage display (ProP-PD)

Wildtype CN and mutants thereof (CN_NIR_ and CN_WF_) were used as bait proteins in selections against a recently described M13 phage library that displays sixteen amino acids peptides tiling intrinsically disordered regions of the human proteome on the major coat protein p8 (Davey et al., 2017) Selections were performed following the published protocol (Davey et al., 2017) with slight modifications. In an attempt to assess the calcium dependence of the interactions, selections were performed in parallel using either 2 mM EDTA or 2 mM CaCl_2_ during incubation of phages with bait proteins and during all washing steps. The presence of calcium is expected to enhance LxVP binding to CN (Rodriguez et al., 2009). For phage selections, 20 μg proteins (GST, CN_WT_, CN_NIR_ and CN_WF_) in 100 μl TBS (20 mM Tris-HCl, 150 mM NaCl, pH 7.4) were immobilized in a 96-well MaxiSorp plate (Nunc, Roskilde, Denmark) overnight at 4°C. Wells were thereafter blocked with 0.5% (w/v) bovine serum albumin (BSA) in TBS (1h, 4°C). Before each round of selection, the phage library (~10^12^ phage virions/100 μl TBT buffer (TBS + 0.05% (v/v) Tween20 + 0.5% (w/v) BSA)) was subjected to a pre-selection toward immobilized GST to remove non-specific binders (1 h, 4°C). The phage library was then transferred to target wells and the phages were allowed to bind for 2 h at 4°C. Target wells were washed five times with cold wash buffer (TBS, 0.5% v/v Tween-20) and retained phages were eluted with log phase (OD_600_ = 0.8) *E. coli* Omnimax in 2xYT (10 g yeast extract, 16 g tryptone, 5 g NaCl to 1 l H_2_O) by incubating for 30 min at 37°C with shaking. After incubation, M13KO7 (NEB, Ipswich, MA, USA) helper phages (1×10^11^ p.f.u./ml) were added to enable phage production and the cells were grown for 45 min at 37°C. To amplify phages, eluted phages were grown over night at 37°C in 10 ml 2xYT containing 100 μg/ml carbenicillin, 30 μg/ml kanamycin, and 0.3 mM IPTG. To determine the number of phages recovered from the selection rounds, dilution series of the infected bacteria were deposited on 2xYT plates containing 100 μg/ml of carbenicillin.

Bacteria were grown overnight and pelleted by centrifugation (4,000 X g, 10 min). Phages were harvested by addition of 2.5 ml ice-cold PEG/NaCl solution (20% w/v PEG 8000 + 2.5 M NaCl) to the supernatant, followed by incubation on ice for 10 min and centrifugation at 10,000 X g for 20 min. Phage pellets were re-suspended in 1 ml TBS and used for the next round of selections. After four rounds of selection, phage pools for each selection round were subjected to enzyme-linked immunosorbent assays as described elsewhere (Huang and Sidhu, 2011).

Enriched phage pools were barcoded for next-generation sequencing. Undiluted amplified phage pools (5 μl) were used as templates for 24 cycles 50 μl PCR reactions using a distinct set of barcoded primers (0.5 μM each primer) for each reaction and Phusion High Fidelity DNA polymerase (NEB) with a maximum polymerase concentration. PCR reactions were supplemented with Gel Loading Dye Purple (6x) (New England Biolabs) and separated on a 2.5 % low melt agarose (BioRad) gel stained with Roti-Safe GelStain (Carl-Roth). The DNA was visualized by UV light. The PCR products were cut out and extracted using the QIAquick Gel Extraction Kit (Qiagen) according to the manufacturer with the following exceptions: a) Gel extracts were resolved at room temperature; b) DNA was eluted with 30 μL low Tris-EDTA (TE) buffer (Life Technologies). Molarities of the eluted library DNA were determined on the 2100 Bioanalyzer using the High Sensitivity DNA Kit (Agilent). Template preparation was performed according to the manufacturer’s instruction using the Ion PGM Template OT2 200 Kit on the Ion OneTouch 2 System (Thermo Fisher Scientific). 25 μL of 5 pM library DNA (1.25*10^-4^ pmol) were used in the template reaction. Sequencing was conducted on the Ion Torrent PGM sequencer using the Ion PGM Sequencing 200 Kit v2 and the Ion 314 Chip v2 (Thermo Fisher Scientific) according to the manuals. Signal processing and base calling were done using the Torrent Suite Software (Thermo Fisher Scientific). Reads were exported as FASTQ files for downstream processing. Phage selections were not successful with CN_WF_ (Fig. S1B). Thus, peptides recovered with this mutant were discarded. Enriched motifs in the sequenced phage pools were identified using SLiMsearch (Krystkowiak and Davey, 2017). Settings, background frequencies, motifs discovered and statistics are reported in Table S1.

### In vitro peptide binding assays

Peptides were fused to GST in vector pGX4T-3 and expressed in *E. coli* BL21 (DE3) (Invitrogen). Cells were grown at 37°C to mid-log phase and induced with 1 mM IPTG for 2 hr. Cell lysates were prepared using the EasyLyse™ bacterial protein extract solution (Lucigen Corp. USA) or the CelLytic B reagent (Sigma, USA) according to the manufacturers’ protocol and were stored at −80°C. 1-2 μg His-tagged calcineurin (truncated and activated to ensure binding to LxVP SLiMs) was first bound to Ni-NTA-agarose or magnetic Dynabeads (Thermo Fisher Sci. USA) in base buffer (50 mM Tris-HCl pH 7.5, 150 mM NaCl, 0.1% Tween 20, 1mM β-mercaptoethanol, protease inhibitors, 5-10 mM imidazole, 1 mg/ml BSA) for 1 hr at 4°C. 50-100 μg of bacterial cell lysate containing GST-peptide was then added to the binding reaction and incubated further for 2-3 hr. 3% of the reaction mix was removed as ‘input’ prior to the incubation, boiled in 2XSDS buffer and stored for downstream analysis. The beads were washed in base buffer containing 15-20 mM imidazole. Bound proteins were then extracted with 2X-SDS loading buffer by boiling for 5 min. The proteins were analyzed by SDS-PAGE and immunoblotting with anti-GST (BioLegend MMS-112P) and anti-His (Qiagen 34660) antibodies. Blots were imaged with the Li-Cor Odyssey imaging system. GST proteins co-purifying with HIS-CN were normalized to their respective input and amount of calcineurin pulled down. Copurification with CN was reported relative to that of a known SLiM (LxVP or PxIxIT) from NFATC1 and GST alone, with positive binders showing significantly more co-purification with CN than GST alone (p<0.01). In each case, 3 or more independent experiments were performed. Statistical significance was determined with unpaired Student’s T test, using GraphPad. For Fig. 1E, NOTCH1 ICD peptides used were NOTCH_WT_: NTPSHQ**L**Q**VP**EHPFLT and NOTCH1_MUT_: NTPSHQ**<u>A</u>**Q**<u>AA</u>**EHPFLT. For Fig. 1G, NUP153 peptdides used were NUP153_WT_: L**P**T**F**N**FS**SPEITTSSP and NUP153_MUT_: L**<u>A</u>**T**<u>A</u>**NF**<u>A</u>**SPEITTSSP.

### Candidate-based BioID Analyses

HEK293T cells were co-transfected with “bait” (Myc-BirA*-CN constructs) and “prey” (FLAG-tagged candidate substrates) using calcium phosphate transfection. Media was changed 16 hours after transfection. 24 hours after transfection, cells were treated with fresh media containing 50 μM D-biotin (Sigma B-4501). After 24 hours of labeling, cells were collected and snap frozen in liquid nitrogen. Cells were lysed in RIPA-2 buffer (150 mM NaCl, 1% NP40, 0.5% Deoxycholate, 0.1% SDS, 50 mM Tris 8.0) on ice for 20 minutes and spun for 20 minutes at 13,000 RPM at 4°C. For each binding reaction, 2 mg of clarified lysate was incubated with 40 μL of rinsed streptavidin beads (Sigma 11641786001) overnight at 4°C. Beads were rinsed and rotated 3 x 10 minutes in RIPA-2 buffer and eluted in 2X sample buffer (12% SDS, 0.06% Bromophenol blue, 60% glycerol, 375 mM Tris-HCl pH 6.8, 0.6M DTT). Input (40 μg) and bound (50% of total binding) samples were resolved by SDS-PAGE and immunoblotted with FLAG (1:2,000; Sigma F1804), Myc (1:3,000; Cell Signaling 2276S), and β-Actin (1:3,000; Li-Cor Biosciences 926-42210) antibodies. Binding was quantified as FLAG/Myc bound signal normalized to FLAG/Actin Input signal. Statistical significance was determined with unpaired Student’s T test, using GraphPad.

### In vivo dephosphorylation of NOTCH1 intracellular domain (NICD)

HEK293 Flp-In T-REx cells expressing WT or LxVP_MUT_ (_2502_LQVP_2505_ è AQAA) NICD-FLAG constructs were transfected with pcDNA3 (vector alone) or HA-tagged CN_trunc_ and induced with 1 μg/mL doxycycline. Media was changed and refreshed with doxycycline 24 hours after transfection. The following day, cells were treated with DMSO or FK506 (1 μM) for 6 hours and then harvested. Cells were lysed with RIPA-2 buffer and run out on 7.5% SDS-PAGE gels followed by rapid transfer on iBlot2 (Thermo Fisher). Changes in electrophoretic mobility were assessed via immunoblotting with FLAG (Sigma F3165), HA (Sigma H3663), and Actin (LiCor 926-42210) antibodies. NICD-FLAG steady-state levels were quantified using ImageStudio imaging software and normalized as FLAG/Actin.

### In vitro dephosphorylation of NICD

HEK293 Flp-ln T-REx cells expressing WT or LxVP_mut_ (_2502_LQVP_2505_ è AQAA) NICD-FLAG constructs were grown in 10-cm dishes and induced with 1 μg/mL doxycycline and collected after 48 hours. Pellets were re-suspended in lysis buffer (50 mM Tris-HCL pH 7.5, 150 mM NaCl, 1% NP-40, 1mM DTT and protease inhibitors) supplemented with phosphatase inhibitors (5 mM NaF, 0.4 mM sodium orthovanadate, 0.1 mM β-glycerophosphate, 5 mM EDTA) by gently vortexing followed by fine-needle aspiration and then incubated on ice for 20 minutes. Lysates were prepared by centrifugation at 15,000g for 20 minutes and removing the supernatant. Total protein concentration was calculated with a BCA protein assay kit (ThermoFisher, USA). Approximately 750 μg of total protein lysate per pull-down was bound to 50 μl of Flag-M2 magnetic beads (Sigma-Aldrich, USA) in binding buffer (lysis buffer but with 1%-NP-40) at 4°C for 2 hr. on an end-over-end rotator. Flag-tagged proteins were pelleted at 2K for 2 min., washed 2X in binding buffer and then once in binding buffer lacking phosphatase inhibitors. The beads were then washed with λ-phosphatase PMP buffer (1X PMP buffer, 1mM MnCl_2_, protease inhibitors), PMP buffer with phosphatase inhibitors (described above), calcineurin buffer (50 mM Tris-HCl, pH 7.5, 100 mM NaCl, 6 mM MgCl_2_, 0.5 mM CaCl_2_) or calcineurin buffer with phosphatase inhibitors, as required. 0.25 μl λ-phosphatase or 100 nM activated calcineurin (Bond et al., 2017) was then added to the reactions followed by incubation at 30°C for 45 min. Reactions were stopped with 2X SDS dye and boiled for 5 min. Proteins were analyzed on 5-5.5 % SDS-PAGE gels followed by Western blotting with the anti-Flag M2 antibody (Sigma-Aldrich, USA).

### PSSM Analyses

#### Dataset construction

A set of experimentally characterized *CN-binding motifs* (CN-BM) for the PxIxIT- and LxVP-binding pockets was curated from (i) the newly validated peptides from the current study and (ii) the previously validated peptides from the CN literature (Table S2). A peptide from NFX1, that was discovered by the ProP-PD screen and validated as binding to the LxVP-pocket (Table S1), was also included as an LxVP motif. The CN-BM dataset contained 33 PxIxIT and 34 LxVP motifs. A background set of *unvalidated consensus matches* (CN-UCM) of PxIxIT- and LxVP-like peptides was constructed by searching the CN-binding PxIxIT- and LxVP motif consensuses (PxIxIT – P[^PG][ILVF][^PG][ILVF][^PG] and LxVP – Lx[VLI]P) against the human UniProt reviewed proteins and removing validated *CN-binding motifs.* The CN-UCM set will consist almost exclusively of peptides that cannot bind to CN but resemble the motifs specificity determinants of the CN motif-binding pocket. The CN-UCM peptide dataset contained 21,791 PxIxIT and 13,014 LxVP consensus matches (Table S2).

#### PSSM construction

Validated CN-binding peptides from the CN-BM set were trimmed, based on position-specific enrichment of amino acids, or physicochemical groupings of amino acids, as described for the consensus representation in the PSSMSearch tool, to create peptide alignments with the consensus xxxP[^PG][ILVF][^PG][ILVF][^PG]xxx for PxIxIT-motifs and xxLx[VLI]Px for the LxVP-motifs (Table S2). A position-specific scoring matrix (PSSM) was constructed from each alignment using the PSI BLAST IC scoring scheme of the PSSMSearch PSSM construction tool (Altschul et al., 1997; Krystkowiak et al., 2018) (Table S2).

#### Discriminatory Attribute Benchmarking

The ability of a range of motif attributes to correctly differentiate *binding* from *background* peptides was tested by analyzing the CN-BM and CN-UCM peptide sets. The classes of motif attributes tested were (i) intrinsic disorder propensity using IUPred and FELLS (Dosztanyi et al., 2005; Piovesan et al., 2017), (ii) disorder-to-order transition upon binding propensity using Anchor (Dosztanyi et al., 2009), (iii) secondary structure propensity using FELLS (helix and extended) and Hydrophobic Cluster Analysis (Piovesan et al., 2017), (iv) conservation using taxonomic range and relative local conservation probability on metazoan alignments (Davey et al., 2012; Krystkowiak and Davey, 2017) and (v) shared molecular function, localization and biological process ontology terms with CN (Krystkowiak and Davey, 2017). The discriminatory ability of the attributes were measured as (i) the AUC *(Area Under The Curve)* of a ROC *(Receiver Operating Characteristics)* curve to determine if the attributes are capable of distinguishing between *binding* and *background* peptides and (ii) Mann-Whitney U test p-values to determine whether the attribute values for the *binding* and *background* peptides were from the same distribution (Table S2). The ability of the PSSM to discriminate *binding* from *background* peptides was tested differently using *leave-one-out* crossvalidation as the PSSM is built using the *binding* peptides. Each peptide in the CN-BM set was removed from the alignment for PSSM construction, the resulting PSSM was used to score the removed peptide and the rank/significance of the removed peptide was calculated (Table S2).

#### Peptide Filtering

The benchmarking analysis identified the PSSM score and accessibility as key discriminatory attributes. Therefore the consensus matches were filtered using the PSSM score *p-value* with a cut-off of 0.0001, localization based on intracellular localization GO terms, and accessibility based on: (i) overlap with a resolved region in a structure from PDB, (ii) intrinsic disorder predictions (retaining only peptides found in disordered regions as defined by an IUPred score ≥0.3 (Dosztanyi et al., 2005) and (iii) UniProt annotation of topologically inaccessible regions (e.g. transmembrane and extracellular regions) (UniProt, 2015). Applying these simple filtering criteria, we retained 409 putative PxIxIT motifs and 410 putative LxVP motifs in the human proteome (Table S2). The predictive power of each discriminatory attribute was recalculated as described in the benchmarking section after filtering was applied showing that no further filtering steps were available to improve confidence in the returned peptides (Table S2). However, relative local conservation probability remained significant in the Mann-Whitney U tests and can be used along with PSSM scores to rank peptides by confidence. This is intuitive as it suggests that peptides that are more conserved and more similar to validated peptides are more likely to be binders.

#### CN interactor enrichment

Distinct CN interaction datasets were created from several sources: the HIPPIE, BioGRID, and IntAct databases, the HIPPIE database filtered by high-throughput methods, BioID data from the PDB-MS study (docking site-dependent and -independent), nonspecific BirA*-labelled proteins (http://www.crapome.org; Workflow 2>Cell/tissue type: HEK293>Epitope tag: BirA*-FLAG>Fractionation: 1D LC-MS>Affinity Approach 1: Streptavidin>Affinity Support 1: Agarose), MAPK docking site-containing proteins and a manually curated set of interactors created for this project (PRS) (Table S5). The enrichment of predicted CN-binding motifs in each interaction dataset was quantified using a randomized PSSM approach. Each PSSM was shuffled 1,000 times, the shuffled PSSMs were searched against the human proteome, and the returned peptides were scored and filtered as described above. The number of CN interactors was calculated for each “shuffled” dataset. The distribution of interactor counts for the 1,000 shuffled PSSM peptide sets was then used to quantify the likelihood of the observed number of interactors with predicted CN-binding peptide-containing proteins (Table S2).

#### Web resource

Variants of the PSSMSearch tool, to analyses proteomes for regions with significant similarity to a specificity determinant model (Krystkowiak and Davey, 2017), and the ProViz tool, to investigate the functional and evolutionary features of a protein (Jehl et al., 2016), allowing PxIxIT and LxVP searches to be performed on the proteome and protein level are available at http://slim.icr.ac.uk/motifs/calcineurin/.

### BioID/MS Analyses

#### Cell line generation

BirA*-FLAG constructs were generated via Gateway cloning into pDEST 5’ BirA*-FLAG pcDNA5 FRT TO. Stable cell lines were generated in HEK293 Flp-In T-REx cell pools as in (Kean et al., 2012). Expression was induced with tetracycline (1μg/ml) for 24 hrs when cells reached 80% confluency on a 150mm plate in selection media. For BioID experiments, cells were treated with 50μM biotin alongside tetracycline induction for 24hrs.

#### Streptavidin affinity purification and data acquisition

Cell pellets were resuspended in a 1:4 ratio of pellet weight to radio-immunoprecipitation assay (RIPA) buffer [50 mM Tris-HCL (pH 7.5), 150 mM NaCl, 1% NP-40, 1 mM EGTA, 4.5 mM MgCl2 and 0.4% SDS), supplemented with 250U/ml of benzonase and 1x Sigma protease inhibitors P8340. Cells were lysed in one freeze-thaw cycle (frozen on dry ice for 5 min, then transferred to 37°C water bath until a small amount of ice remained). The sample was gently agitated on a nutator at 4°C for 30 min and then centrifuged at 16,000g for 20 min at 4°C. The supernatant was transferred to 1.5 ml microcentrifuge tubes, and a 25 μl volume of 60% streptavidin-sepharose bead slurry (GE Healthcare, catalog no. 17-5113-01) was added. Prior to addition of beads to supernatant, the beads were washed three times with RIPA buffer and resuspended in RIPA buffer to make a 60% slurry. Affinity purification was performed at 4°C on a nutator overnight, beads were then washed once with 2% SDS buffer (500 μl of 2%SDS, 50 mM Tris pH 7.5), twice with 500 μl of RIPA buffer and three times with 50 mM ammonium bicarbonate pH 8.0 (ABC). After removal of the last wash, the beads were resuspended in 100 μl of 50 mM ABC (pH 8.0) with 1 μg of trypsin (Sigma-Aldrich, T6567) and rotated on a nutator at 37°C for 4hrs. After 4 hr., an additional 1 μg of trypsin was added to each sample (in 2 μl of 50 mM ABC) and rotated on a nutator at 37°C overnight. Beads were pelleted (500 g, 1 min) and the supernatant (pooled peptides) was transferred to a new 1.5-ml microcentrifuge tube. Beads were rinsed with 100 μl of MS-grade H_2_O and combined with the pooled peptides. 50 μl of 10% formic acid was added to the supernatant for a final concentration of 2% formic acid. The pooled supernatant was centrifuged at 16,000g for 5 min to pellet remaining beads. 230 μl of the pooled supernatant was transferred to a new 1.5-ml microcentrifuge tube. Samples were dried using a vacuum concentrator. Tryptic peptides were resuspended in 10 μl of 5% formic acid, 2.5 μl was used for each analysis.

Peptides were analyzed by nano-HPLC (high-pressure liquid chromatography) coupled to MS. Nano-spray emitters were generated from fused silica capillary tubing (100μm internal diameter, 365 μm outer diameter) using a laser puller (Sutter Instrument Co., model P-2000, heat = 280, FIL = 0, VEL = 18, DEL = 2000). Nano-spray emitters were packed with C18 reversed-phase material (Reprosil-Pur 120 C18-AQ, 3 μm) in methanol using a pressure injection cell. 5ul of sample (2.5 μl of each sample with 3.5 μl of 5% formic acid) was directly loaded at 800 nl/min for 20 min. onto a 100 μm x15 cm nano-spray emitter. Peptides were eluted from the column with an acetonitrile gradient generated by an Eksigent ekspert™ nanoLC 425, and analyzed on a TripleTOF™6600 instrument (AB SCIEX, Concord, Ontario, Canada). The gradient was delivered at 400 nl/min from 2% acetonitrile with 0.1% formic acid to 35% acetonitrile with 0.1% formic acid using a linear gradient of 90 min. This was followed by a 15min wash with 80% acetonitrile with 0.1% formic acid and equilibration for another 15min to 2% acetonitrile with 0.1% formic acid, resulting in a total of 120min for the DDA (data-dependent acquisition) protocol.

The first MS1 scan had an accumulation time of 250 ms within a mass range of 400-1800Da. This was followed by 10 MS/MS scans of the top 10 peptides identified in the first DDA scan, with an accumulation time of 100ms for each MS/MS scan. Each candidate ion was required to have a charge state from 2-5 and a minimum threshold of 300 counts per second, isolated using a window of 50mDa. Previously analyzed candidate ions were dynamically excluded for 7 seconds. The DIA (data-independent acquisition; SWATH) protocol used the same gradient and nanoLC protocol as the DDA method. The MS1 scan had an accumulation time of 250ms within a mass range of 400-1800Da, followed by MS/MS scans split into 54 scan windows across the mass range of 400-1250Da. In each MS/MS scan window, the accumulation time was 65ms per MS/MS scan.

Both DDA and SWATH datasets has been deposited as a complete submission to the MassIVE repository (https://massive.ucsd.edu/ProteoSAFe/static/massive.jsp). The DDA data has been assigned the accession number MSV000083695. The ProteomeXchange accession is PXD013527. The dataset is currently available for reviewers at ftp://MSV000083695@massive.ucsd.edu. Please login with username MSV000083695_reviewer; password: Calcineurin2019. The SWATH data has been assigned the accession number MSV000083697. The ProteomeXchange accession is PXD013529. The dataset is currently available for reviewers at ftp://MSV000083697@massive.ucsd.edu. Please login with username MSV000083697_reviewer; password: CalcineurinS2019. Both datasets will be made public upon acceptance of the manuscript.

#### MS data analysis

Mass spectrometry data generated were stored, searched and analyzed using ProHits laboratory information management system (LIMS) platform (Liu et al., 2016). Within ProHits, WIFF files were converted to an MGF format using the WIFF2MGF converter and to a mzML format using ProteoWizard (V3.0.10702) and the AB SCIEX MS Data Converter (V1.3 beta). For DDA, the data was then searched using Mascot (V2.3.02) (Perkins et al., 1999) and Comet (V2016.01 rev.2) (Eng et al., 2013). The spectra were searched with the human and adenovirus sequences in the RefSeq database (version 57, January 30^th^, 2013) acquired from NCBI, supplemented with “common contaminants” from the Max Planck Institute (http://maxquant.org/contaminants.zip) and the Global Proteome Machine (GPM; ftp://ftp.thegpm.org/fasta/cRAP/crap.fasta), forward and reverse sequences (labeled “gi|9999” or “DECOY”), sequence tags (BirA, GST26, mCherry and GFP) and streptavidin, for a total of 72,481 entries. Database parameters were set to search for tryptic cleavages, allowing up to 2 missed cleavages sites per peptide with a mass tolerance of 35 ppm for precursors with charges of 2+ to 4+ and a tolerance of 0.15amu for fragment ions. Variable modifications were selected for deamidated asparagine and glutamine and oxidized methionine. Results from each search engine were analyzed through TPP (the Trans-Proteomic Pipeline, v.4.7 POLAR VORTEX rev 1) via the iProphet pipeline (Shteynberg et al., 2011).

For DiA, MSPLIT-DIA v1.0 (Wang et al., 2015) was used to analyze the data within the ProHits-LIMS platform (Wang et al., 2015). A unique peptide-spectrum library was generated from the peptide-spectrum matches (PSMs) from the matched DDA runs (30 runs). Only those with the lowest MS-GFDB (Kim et al., 2010) probability, with respect to each unique peptide sequence and precursor charge state, and passed a peptide level false-discovery rate (FDR) of 1% using the Target-Decoy approach (Elias and Gygi, 2007). The MS-GFDB was set to search for tryptic cleavages with no missed cleavage sites. In addition, the peptide length was restricted from 8 to 30 amino acids and required precursors to be within a mass tolerance of 50ppm and have charge states 2+ to 4+. Fragment ions were searched with a tolerance of +/-50ppm. Oxidized methionine was selected as a variable modification in the MS-GFDB search. The NCBI RefSeq database (version 57, January 30^th^, 2013) was used to search the spectra. This database contains 36241 human and adenovirus sequences and was supplemented with common contaminants from Max Planck Institute (http://maxquant.org/contaminants.zip) and the Global Proteome Machine (GPM; ftp://ftp.thegpm.org/fasta/cRAP/crap.fasta). The spectra library also incorporates non-redundant PSMs from the SWATH-Atlas library (https://www.nature.com/articles/sdata201431). Additional decoys were added to the spectra library using the decoy library command built in to MSPLIT with a fragment mass tolerance of +/−0.05 Da. This spectral library was used for protein identification of DIA data using MSPLIT (Wang et al., 2015). The MSPLIT search required parent mass tolerance of +/− 25 Da and a fragment mass tolerance of +/− 50ppm.

### SAINT analysis

SAINTexpress version 3.6.1 was used as a statistical tool to calculate the probability of potential protein-protein interactions from background contaminants using default parameters (Teo et al., 2014). SAINT analysis was performed using two biological replicates per bait for both DDA and SWATH. Twenty-two negative control experiments were conducted for BioID; eleven with 3xFLAG-alone samples and eleven with BirA*FLAG-alone samples. Controls were compressed to 11 samples and two unique peptide ions and a minimum iProphet probability of 0.99 were required for protein identification. SAINT probabilities were calculated independently for each sample, averaged (AvgP) across biological replicates and reported as the final SAINT score. Only SAINT scores with a FDR ≤ 1% were considered high-confidence protein interactions.

### Immunofluorescence of centrosomal proteins

HEK293 Flp-ln T-REx cells were grown on poly-L-lysine-coated #1.5 glass coverslips (Electron Microscopy Sciences). Cells were treated with 1 μg/mL doxycycline for 24 h to induce expression of BirA* or BirA*-CNAα. Media was then replaced with fresh media containing 1 μg/mL doxycycline and 50 μM biotin, and labeling was allowed to proceed for 18 hr. Cells were fixed with −20°C methanol for 15 minutes, washed with PBS and blocked with PBS-BT (3% BSA, 0.1% Triton X-100, 0.02% sodium azide in PBS) for 30 min. Coverslips were incubated with primary antibodies diluted in PBS-BT for 1 hr., washed with PBS-BT, incubated with secondary antibodies and DAPI diluted in PBS-BT for 1 hr., then washed again. Samples were mounted using Mowiol (Polysciences) in glycerol containing 1,4,-diazobicycli-[2.2.2]octane (DABCO, Sigma-Aldrich) antifade.

Primary antibodies used for immunofluorescence: mouse IgG2b anti-centrin3, clone 3e6 (1:1000, Novus Biological) and mouse IgG1 anti-gamma-tubulin, clone GTU-88 (1:1000, Sigma-Aldrich). For detecting biotinylation: Alexa594-streptavidin (1:1000, Thermo Fisher). Secondary antibodies used for immunofluorescence were diluted 1:1000: Alexa647 anti mouse IgG1 (ThermoFisher) and Alexa488 anti mouse IgG2b (ThermoFisher). For Fig. 4E, images were acquired as 0.3695 μm Z-stacks on a Leica SP8 scanning laser confocal microscope with a 63x/1.4 NA oil objective, using white light laser excitation at 80% power, and LASX Software. For Fig. S5C and D, images were acquired as 0.25 μm Z-stacks using a Zeiss Axio Observer microscope with a confocal spinning-disk head (Yokogawa), PlanApoChromat 63x/1.4 NA oil objective, and a Cascade II:512 EM-CCD camera (Photometrics), run with MicroManager software (Edelstein et al., 2014). Post-imaging processing was performed using Fiji (Schindelin et al., 2012). Images are maximum projections of confocal stacks.

### Proximity Ligation Assays between CN and nuclear basket nucleoporins

To analyze interactions between endogenous CN and nuclear basket nups *in vivo* (Figure 5B and Supplemental Figure 5A), HeLa cells were seeded onto coverslips and subjected to Proximity Ligation Assays (PLAs) 24 hours later using the Duolink RED proximity ligation assay kit (Sigma; DUO92101). PLAs were carried out according to manufacturer’s instructions with primary antibodies against CN (Rabbit, Bethyl 908A, 1:250), NUP50 (Mouse, SantaCruz G-4, 1:500), NUP153 (Mouse, Abcam ab96462, 1:250), and TPR (Mouse, SantaCruz H-8, 1:50). All samples were imaged on a single z-plane on the Lionheart™ FX automated widefield microscope with a 20X Plan Fluorite WD 6.6 NP 0.45 objective. DAPI (365 nm), GFP (465 nm), and Texas Red (590 nm) filter cubes were used to visualize nuclei, GFP-peptide expression (see next section), and PLA puncta, respectively. All fields within each biological replicate were captured using the same channel exposure and gain to ensure representative trends across treatments. The number of nuclear PLA puncta/cell was calculated in FIJI (macros: https://github.com/cyertlab/CNome.git) for a range of 421 to 529 cells between conditions and across 2 biological replicates.

For GFP-peptide analyses (Figure 5C and Supplemental Figures 5B-D), 7.5 x 10^5^ HeLa cells were seeded onto coverslips in a 60-mm dish and transfected with 4 μg of plasmid DNA using JetOPTIMUS transfection reagent (VWR). 24 hours after transfection, cells were subjected to PLA analyses using the same reagents, antibody dilutions, imaging approach, and analysis described above. Due to very high transfection efficiency and GFP-peptide expression, all cells were considered for nuclear PLA puncta analyses. GFP-peptide PLA analyses were performed in triplicate and the number of cells across conditions ranged from 1,332 to 1,605. Statistical analyses comparing nuclear PLA puncta between GFP-peptide samples were carried out using the Kolmogorov-Smirnov test to determine significance (p-value) and maximum difference between distributions (D statistic) using GraphPad PRISM.

### In vitro interaction between CN and human nucleoporins

WT and PxIxIT mutant (P.M.) versions of GST-tagged Nup153_228-611_ (P.M. = _485_PTFNFS_490_ ⇒ ATANAA), TPR_626-2363_ (P.M. = _2083_PRLTIH_2088_ ⇒ ARATAA), and Nup50 (full length; P.M. = _345_PKVVVT_350_ ⇒ AKAVAA) were purified from BL21-DE3 (Stratagene 230245) cells as described above for GST-tagged CN. For binding reactions, purified His-tagged CN (purification described above) was incubated with ~1 μg GST-NUPs (WT and PxIxIT_MUT_) or GST alone at 4°C for 1 hour with rotation (base binding buffer: 50 mM Tris-HCl pH 7.5, 150 mM NaCl, 0.1% Triton-X 100, 10 mM Imidazole, 1 mM BME and protease inhibitors). Following GST-His incubation, equilibrated Ni-NTA beads (Fisher PI25214) were added to each reaction and incubated for another 2 hours at 4°C with rotation. Beads were washed three times in base binding buffer supplemented with 15 mM Imidazole, followed by elution with 2X sample buffer at 37°C. Input (3% of total binding reaction) and bound samples (50% of binding reaction) were subjected to SDS-PAGE followed by immunoblot analysis with His (Qiagen 34660) and GST (BioLegend MMS-112P) antibodies.

### In vitro dephosphorylation of human nucleoporins

Constitutively active His-tagged ERK2 (Addgene # 39212) was purified from BL21-DE3 RIPL cells (Stratagene 230280) as previously described (Khokhlatchev et al., 1997). WT and PxIxIT mutant (P.M.) versions of GST-tagged Nup153_228-611_ (P.M. = _485_PTFNFS_490_ ⇒ ATANAA), TPR_626-2363_ (P.M. = _2083_PRLTIH_2088_ ⇒ ARATAA), and Nup50 (full length; P.M. = _345_PKVVVT_350_ ⇒ AKAVAA) were purified from BL21-DE3 (Stratagene 230245) cells as described above for GST-tagged CN.

For phosphorylation/dephosphorylation reactions, first 1-10 μg of purified GST-NUPs were phosphorylated with 50 ng purified ERK2 in kinase buffer (20 mM Tris-HCL pH 7.5, 15 mM MgCl_2_, 1 mM EGTA, 2mM DTT and phosphatase inhibitors) supplemented with 100 μM ATP and γ^32^-P ATP (5 μCi/reaction) for 12-16 hr. at room temperature (RT). Unincorporated ATP was removed using a Centri-Sep spin column (ThremoFisher 401762). In order to remove ERK2, the reactions were incubated with 10 μl of rinsed Ni-NTA beads for 1hr. at RT, spun briefly and supernatant transferred to new tubes. Reactions were then supplemented with 1mM CaCl_2_, 100 nM purified His-CN (purified as described above) and 200 μM competing peptides (VIVIT or SCR) and incubated at 30°C. Time points were collected at 0, 5, 10 and 30 minutes after addition of CN and the reactions stopped by heating with 2X SDS dye. Samples were subjected to SDS-PAGE analysis followed by Gel Code staining (Fisher P124590), then dried in a gel drier and exposed to a phosphor-imager screen to quantify incorporated ^32^-P signal. ^32-^P signal was normalized to protein amount. The data represent an average of 3 experiments. Statistical significance was determined with unpaired Student’s T test, using GraphPad PRISM.

### Time Lapse Imaging and Nuclear Import Assay

HeLa cells grown in DMEM supplemented with 10% FBS were plated on 25 mm glass coverslips and transfected (Lipofectamine LTX; Life Technologies) with a plasmid encoding a chimeric Rev-GFP-Glucocorticoid Receptor protein [RGG; (Love et al., 1998)], whose import into the nucleus is induced by addition of dexamethasone. Twenty-four hours after plasmid transfection, cells were pretreated for 1 hour with 1 μM FK506, 2 μM cyclosporin A, or an equivalent volume of DMSO (vehicle control). Coverslips of cells were subsequently mounted in a Ludin Chamber (Life Imaging Services) and changed to DMEM/F12 supplemented with 10% FBS and HEPES buffer. Nuclear import of the RGG biosensor was induced in the presence of DMSO, FK506, or cyclosporin A by adding dexamethasone at a final concentration of 100 nM. Imaging was performed on an inverted Nikon T*i*-E microscope (Nikon Instruments) equipped with a Nikon 20X Plan Apo NA 0.75 objective. This system utilized an Andor spinning disk confocal (Andor Technology) and an Andor Ixon 885 EMCCD camera. Time lapse sequences were captured at 30 second intervals for 90 minutes using the Andor IQ imaging software. Regions of interest containing the nuclear and cytoplasmic compartments of individual cells were manually drawn and applied to all images in the time-lapse sequence stack. Fluorescence intensity in these regions was measured using the ImageJ software (NIH) and graphed as a ratio of nuclear/cytoplasmic fluorescence for each cell. Initial import rates (Figure 6B) were determined by linear regression analysis of RGG N:C ratios.

### Phosphoproteomic Analysis of Nuclear Pore Complex Proteins

#### Cell treatment and collection

For each biological replicate (total of 3), ~3.6 x 10^7^ HeLa cells were treated with 2 μM Cyclosporin A (CsA; Sigma-Aldrich) or an equivalent volume of DMSO (vehicle control) for 1 hour. After drug treatment, cells were collected on ice with scraping, rinsed in ice-cold PBS, pelleted by centrifugation, and snap frozen in liquid nitrogen.

#### Cell lysis and Immunoprecipitation

Cell pellets were lysed in RIPA-2 buffer (150 mM NaCl, 1% NP-40, 0.5% deoxycholate, 0.1% SDS, and 50 mM Tris, pH 8.0) supplemented with a protease and phosphatase inhibitor cocktail (Halt^™^, ThermoFisher) and subjected to fine needle aspiration with a sterile 27.5 gauge needle. Cell lysates were clarified by centrifugation (13,000 RPM for 20 minutes) and subjected to BCA analysis to measure protein concentration. For each sample, 4 mg of clarified lysate was incubated with antibody-coated protein G beads (3 mg protein G Dynabeads^™^ [ThermoFisher] + 24 μg mAb414 antibody [BioLegend]) in IP buffer (50 mM Tris, pH 7.5, 150 mM NaCl, 0.5% NP-40) supplemented with a protease and phosphatase inhibitor cocktail. IP reactions were carried out for 3 hours at 4°C with end-over-end rotation, followed by 3 x 1 mL rinses with supplemented IP buffer. Bound proteins were eluted from the beads by boiling in 2X sample buffer (4X LDS stock; Invitrogen) and separated by SDS-PAGE on 4-12% Bis-Tris NuPAGE gels (Invitrogen).

#### Peptide generation and analysis

For each lane, an ~1.5 cm area between the 50 kDa IgG band and loading well was excised and cut into 2 segments for in-gel Tryptic digestion. Peptides were desalted using C18 ZipTips and analyzed by liquid chromatography-tandem mass spectrometry (LC-MS/MS) on an Easy LC1200 UPLC liquid chromatography system connected to a Q-Exactive HF hybrid quadrupole-Orbitrap mass spectrometer (Thermo Fisher). Peptides were separated using analytical column ES803A (Thermo Fisher). The flow rate was 300 nl/min and a 120-min gradient was used. Peptides were eluted by a gradient from 3 to 28 % solvent B (80 % acetonitrile, 0.1 % formic acid) over 100 minutes and from 28 to 44 % solvent B over 20 minutes, followed by short wash at 90 % solvent B. Precursor scan was from mass-to-charge ratio (m/z) 375 to 1,600 and top 15 most intense multiply charged precursors were selection for fragmentation. Peptides were fragmented with higher-energy collision dissociation (HCD) with normalized collision energy (NCE) 27. Raw data, search results, and quantification analysis were deposited to PRIDE with identifier number PXD017868; reviewer account user ID: and password:pWWh3kXj.

#### Data processing

Tandem mass spectrometry peak lists were extracted using an in-house script PAVA. Protein Identification and post-translational modification search were done using Protein Prospector (version 5.20.0) (Chalkley et al., 2008) against the random concatenated SwissProt database Homo Sapiens (total 20203 entries). A precursor mass tolerance of 10 ppm and a fragment mass error tolerance of 20 ppm were allowed. Carbamidomethylcysteine was searched as a fixed modification. Variable modifications include protein N-terminal acetylation, peptide N-terminal Gln conversion to pyroglutamate, and Met oxidation. Subsequent searches were performed to find those peptides modified by phosphorylation. The search parameters were as above, but this time allowing for phosphorylation on Ser, Thr and Tyr. Assignments of all modified peptides were checked manually and selected spectra are illustrated in Figure SF6.

Phosphorylation was quantified by using Skyline. Briefly, non-modified and modified peptide MS1 spectra were extracted by Skyline using centroid data with 5 ppm allowance. The peaks were manually curated based on retention time and total area of extracted ions (M, M+1, M+2) of respective peptides were exported to excel file. The calculation was similar as described in (Ni et al., 2013). A value termed the phospho-ratio was defined as a relative level of phosphorylated and non-phosphorylated peptide containing the site. The final ratio is calculated by dividing Phos-Ratio from CsA treated sampled to the one from DMSO control. If CsA has no effect, we would expect to see a final ratio of 1.

In some cases, because the phosphorylation may affect the tryptic cleavage of the peptides, the corresponding non-phosphorylated peptides were not detected. Then the normalization was done by using the total peptide intensities from that protein calculated by Maxquant (version 1.6.2.6). Briefly, settings were default settings. Methionine oxidation and N-terminal acetylation were set as variable modifications and Carbamidomethylcysteine as fixed modification. Maximum number of modifications per peptide was five. Trypsin/P with a maximum of two missed cleavages was set as digest. Peptides were searched against the the same database as mentioned as above plus a list of likely contaminants containing Trypsin, human Keratin and against the contaminants list of MaxQuant. Minimum allowed peptide length was seven. FTMS and TOF MS/MS tolerance were 20 ppm and 40 ppm, respectively, and the peptide FDR and protein FDR were 0.01. Unique and razor peptides were used for protein quantification. LFQ minimum ratio count was set to one and fast LFQ was active with a minimum and average number of neighbors of three and six, respectively. Match between runs and second peptides were checked.

### In vivo interaction between CN and yeast nucleoporins

*GAL1* inducible and N-terminally GST-tagged clones of NUP1 (pEGH-NUP1), NUP2 (pEGH-NUP2) and NUP60 (pEGH-NUP60) were obtained from the yeast GST fusion collection (Zhu et al., 2001). These plasmids were transformed into yeast strain JRY19, a derivative of strain JRY11 (Goldman et al., 2014) containing *3X-GFP-CNA1* and *ACT1-GEV* (Mclsaac et al., 2011). Transformants were grown to mid-log phase when the *GAL1* promoter was induced with 2.5 μM β-estradiol for 4 hr. Cells were treated with 1M NaCl (osmotic stress) and 200 mM CaCl_2_ (to activate calcineurin) for 15 min. prior to harvesting and cell pellets were frozen at −80°C. Pellets were thawed in lysis buffer (50 mM Tris-HCl pH 7.5, 150 mM NaCl, 1% Tween 20, 1mM DTT, protease inhibitors) and lysed with silica beads in a FastPrep-24 homogenizer (MP Biomedicals) with two 40 sec pulses at 6.5 m/sec. Lysed cells were clarified by centrifugation at 20,000g for 20 min. 5-6 mg of lysate was used for pull-downs with 30 μl of GFP-Trap magnetic beads (Bulldog Bio. Inc.) in binding buffer (50 mM Tris-HCl pH 7.5, 150 mM NaCl, 0.1% Tween 20, 1mM DTT, protease inhibitors) containing 200 μM peptides (VIVIT: MAGPHP**VIVIT**GPHEE, or a scrambled version SCR: MAGIVPIHVTHAPgEe) (Aramburu et al., 1999). The binding reactions were incubated at 4°C for 2-4 hr. The beads were washed twice in binding buffer (the first wash containing 200 μM peptides) and co-purifying proteins were extracted with heating in 2X -SDS dye at 37°C for 10 min. Extracted proteins were analyzed by SDS-PAGE followed by Western analysis. GST-proteins were observed with a mouse anti-GST monoclonal antibody and GFP tagged CNA1 was observed with a rabbit anti-GFP polyclonal antibody followed by secondary Li-Cor antibodies. Blots were imaged with the Li-Cor Odyssey imaging system.

### In vitro dephosphorylation of yeast nucleoporins

#### Purification of Hog1 kinase from yeast

Yeast strain BY4741 was transformed with plasmids BG1805-Hog1 and BJ1805-Hog1-KD expressing ZZ-3C-HA-6XHIS tagged, wild-type and kinase-dead mutant versions of Hog1 respectively, from the inducible *GAL1* promoter (a kind gift from Jeremy Thorner). Transformants were grown in 4% raffinose containing medium to mid-log phase followed by induction of the GAL1 promoter with 2% galactose for 20 hrs. at 30°C. Cells were treated with 1M NaCl for 20 min. to activate the Hog1 pathway prior to harvesting. Cells were pelleted, washed with water and frozen. Cell pellets were thawed in lysis buffer (50 mM Tris-HCl pH 7.5, 150 mM NaCl, 0.1% Tween 20, 1mM β-mercapto ethanol, 5mM imidazole with protease and phosphatase inhibitors, as above) and lysed with silica beads in the FastPrep-24 homogenizer (MP Biomedicals) with two 40 sec pulses at 6.5 m/sec. Lysed cells were clarified by centrifugation at 20,000g for 20 min. 60 mg of total cell lysate was bound to 1 ml Ni-NTA beads in lysis buffer at 4°C for 4 hr. Beads were washed with buffer containing 10 mM Imidazole and eluted with 250 mM imidazole. Pooled eluted fractions were dialyzed in buffer containing 50 mM Tris-HCL pH 7.5, 150 mM NaCl, 1mM β-mercapto ethanol.

#### Purification of GST-tagged yeast NUP proteins

Plasmids expressing GST-tagged Nup1 (PSR19), Nup2 (PSR3), and Nup60 (PSR21) in pGEX4T-3 (kind gifts from Francesc Posas) were expressed in BL21 bacterial cells. Cells were grown to OD_600_ 0.7-0.8 and GST-tagged proteins were expressed at 18°C for 16-18 hrs. and purified using glutathione-sepharose beads as detailed above for GST-CN purification.

#### *In vitro* de-phosphorylation of GST-NUPs

~5-10 μg of purified GST-Nup1/2/60 were phosphorylated with 1 μg Hog1 kinase in kinase buffer (50 mM Tris-HCL pH 7.5, 10 mM MgCl_2_, 1mM DTT) supplemented with 100 μM ATP and 2 μl γ^32^-P ATP (3000Ci/mmol) for 2 hr. at 30°C. Unincorporated ATP was removed using a Centri-Sep spin column (ThermoFisher USA). The reactions were then incubated for 1 hr. with 5 μl washed Ni-NTA beads and spun briefly to remove His-Hog1. Supernatants were then incubated with 100nM purified, activated yeast CN (Goldman et al., 2014) that had been pre-incubated with 100μM VIVIT or SCR (see above) for 5 min. at 30°C. De-phosphorylation reactions were carried out at 30°C. Aliquots were removed at 0, 10, 30 and 80 min and the reactions stopped by heating with 2X SDS dye. Samples were subjected to SDS-PAGE. Gels were first stained with Gel Code protein stain, then dried in a gel drier and exposed to a phosphor-imager screen to quantify incorporated ^32^-P signal. ^32-^P signal was normalized to protein amount. The data represent an average of at least 3 experiments. Statistical significance was determined with unpaired Student’s T test, using GraphPad.

